# ModCRE-NN: Interpretable Deep Learning Harnesses Structural and Evolutionary Synergy to Predict Transcription Factor Binding Specificity

**DOI:** 10.64898/2026.05.27.728137

**Authors:** Victor Méndez-Riosalido, Patrick Gohl, Patricia M. Bota, Eric Kramer, Alberto Meseguer, Oriol Gallego, Narcis Fernandez-Fuentes, Baldo Oliva

**Author notes:** To whom correspondence should be addressed. Availability: Contact.

## Abstract

We present ModCRE-NN, a machine-learning framework and server for predicting transcription-factor (TF) DNA-binding motifs through the integration of structural and evolutionary information. The method combines structure-derived Position Weight Matrices (PWMs) together with PWMs of homologous spanning multiple evolutionary sequence-identity intervals, which are integrated into a unified 20-channel tensor representation. Benchmark datasets were constructed on experimental databases of TF motifs, showing DNA binding specificity, while redundancy reduction and strict train/test partitioning minimized homology leakage. Prediction quality was evaluated on an independent separated set of TFs using the similarity analysis of profiles. Three complementary architectures were implemented and evaluated: an interpretable regression-based model, a convolutional neural network (CNN), and a Transformer-based architecture using self-attention mechanisms. The regression model achieved strong performance in high-homology regimes dominated by closely related PWMs, whereas CNN and Transformer architectures showed superior robustness under low evolutionary similarity and increased structural uncertainty. Importantly, AI-generated motifs consistently improved the similarity-scores while reducing prediction variance relative to the original structural and evolutionary input motifs, indicating that the models effectively denoise heterogeneous motif assemblies and reconstruct stable consensus DNA-binding representations rather than simply transferring PWMs from the nearest homolog. The CNN model exhibited the most balanced attribution profile, suggesting enhanced ability to combine weak structural and evolutionary signals into coherent motif representations. Additionally, we implemented a prediction-reliability framework combining Random Forest regression, exponential interpolation, and hybrid residual-corrected modeling to estimate the quality and uncertainty of the PWMs as functions of evolutionary similarity, motif-cluster consistency, and TF-family context. Overall, our results demonstrate that integrating structural information with deep learning provides a robust framework for large-scale TF-binding specificity prediction under conditions of substantial evolutionary divergence and motif uncertainty.

## INTRODUCTION

Transcription factors (TFs) are central regulators of gene expression that control cellular differentiation, tissue homeostasis, responses to environmental stimuli, and disease-associated transcriptional programs [1]. Their regulatory activity depends on the recognition of short DNA sequence motifs located within promoters, enhancers, and other cis-regulatory elements. Consequently, identifying transcription factor binding sites (TFBSs) and understanding the molecular determinants of TF–DNA specificity remain major challenges in molecular biology, genomics, and computational biology [2].

The DNA-binding preferences of TFs have traditionally been represented using Position Weight Matrices (PWMs) or Position Probability Matrices (PPMs), which approximate binding specificity by assuming independent nucleotide contributions at each position of the binding site [3, 4]. Although PWMs remain the dominant representation for TF specificity because of their interpretability and computational efficiency, they only partially capture the complexity of protein–DNA recognition. TF binding is influenced not only by direct amino acid–base interactions (“base readout”) but also by sequence-dependent DNA structural properties (“shape readout”), cooperative interactions among TFs, chromatin accessibility, and contextual flanking sequences [5, 6].

Over the last two decades, large-scale experimental efforts have generated extensive collections of TF-binding profiles using both in vitro and in vivo techniques, including SELEX [7], protein-binding microarrays (PBMs) [8], SMiLE-seq [9], MPRA [10], ChIP-seq [11], ChIP-exo [12], ChIP-nexus [13], and related high-throughput technologies [14]. These approaches have considerably expanded our understanding of TF specificity and regulatory networks. Nevertheless, despite the growing amount of available data, binding specificities remain unknown or poorly characterized for a substantial fraction of eukaryotic TFs [15]. Furthermore, experimental approaches are labor-intensive, expensive, and often highly dependent on specific experimental conditions or antibody quality [16].

To overcome these limitations, computational approaches for TFBS prediction and PWM reconstruction have become increasingly important. One of the most widely used strategies is evolutionary inference, often referred to as the “nearest-neighbor” approach, which transfers binding preferences between homologous TFs sharing similar DNA-binding domains (DBDs) [17]. Databases such as Cis-BP have demonstrated that TFs with highly conserved DBDs frequently retain similar DNA-binding specificities [18–20]. However, evolutionary transfer becomes unreliable at lower sequence identities, particularly within the “twilight zone” of remote homology, where even subtle amino acid substitutions can dramatically alter DNA-recognition preferences [21, 22].

Complementary to evolutionary approaches, structure-based methods attempt to infer DNA-binding specificity directly from the physicochemical properties of protein–DNA interfaces [23, 24]. These methods exploit information derived from experimentally solved or computationally modeled TF–DNA complexes to evaluate the mechanisms governing molecular recognition[25]. Statistical knowledge-based potentials[26] and energy-based scoring functions[27] have been successfully applied to characterize protein–DNA interactions and infer TF-binding preferences from three-dimensional structures[28]. In this context, our previous framework, ModCRE[23], integrated structural modeling with statistical potentials derived from experimental interaction data to predict TF–DNA interactions and reconstruct PWMs even for TFs lacking close homologous motifs[29].

The recent revolution in protein structure prediction driven by RoseTTAFoldNA[30], AlphaFold3[31], Boltz2[32] and other AI-based structural predictors [33], has profoundly transformed the field of structural bioinformatics. These methods now enable the generation of accurate protein and protein–DNA structural models at unprecedented scale, providing a rapidly expanding source of structural information that can potentially be exploited to infer TF-binding specificity. At the same time, advances in artificial intelligence and deep learning have introduced new opportunities to integrate heterogeneous biological information across sequence, structure, and experimental domains.

Recent studies have significantly advanced the field by moving beyond classical motif-transfer approaches toward interpretable and geometry-aware AI frameworks: Bussemaker and colleagues recently developed FamilyCode, a biophysically interpretable machine-learning framework to predict how missense alter TF specificity[21]. In parallel, Rohs and colleagues introduced DeepPBS[24], a geometric deep-learning framework that predicts DNA-binding specificity directly from protein–DNA structural complexes, and Deep DNAshape[34], a deep-learning framework that models long-range DNA-shape dependencies. These studies demonstrated that extended flanking sequences substantially influence DNA structural properties and TF recognition mechanisms, emphasizing the importance of shape readout and long-range context in TF–DNA interactions. Hence, TF specificity prediction is increasingly moving from simple motif transfer toward integrative AI frameworks capable of combining structural, evolutionary, energetic, and contextual information into unified predictive models[16].

In this work, we build upon these developments and propose a machine-learning framework for PWM reconstruction that integrates heterogeneous structural and evolutionary evidence into a unified prediction strategy. Unlike previous approaches relying exclusively on nearest-neighbor transfer or isolated structural analysis, our framework combines structure-derived PWMs generated with ModCRE together with evolutionary PWMs distributed across multiple sequence-identity intervals. These heterogeneous sources are integrated into a 20-channel tensor representation processed through three complementary architectures: an interpretable regression-based model, a convolutional neural network (CNN), and a Transformer-based architecture using self-attention mechanisms.

To increase robustness against motif uncertainty, we implemented a biologically grounded motif-clustering and aggregation strategy combined with stochastic tensor generation and sliding-window decomposition. Additionally, we developed occupancy-corrected attribution analyses and prediction-reliability estimation frameworks that enable biological interpretation of the learned representations and confidence-aware PWM prediction.

Our results demonstrate that deep-learning architectures integrating structural and evolutionary information can reconstruct biologically meaningful TF-binding motifs even under conditions of substantial evolutionary divergence and structural uncertainty. Beyond motif reconstruction itself, the proposed framework provides a general strategy for integrating heterogeneous biological evidence and extracting robust consensus representations from noisy molecular data, offering new opportunities for large-scale characterization of transcriptional regulatory specificity.

## METHODS

### 1. Benchmark construction and evolutionary mapping

Transcription factor (TF) sequences and experimentally derived Position Frequency/Weight Matrices (PFMs and PWMs) were collected from the JASPAR [35], HOCOMOCO [36], and Cis-BP [37] databases. Structural TF monomer and dimer models were generated using the ModCRElib [29], and DNA-binding domain (DBD) regions were extracted for downstream analyses. TFs were additionally classified into evolutionary families using Pfam [38] annotations and HMMER [39].

To minimize homology leakage between training and evaluation datasets, redundancy reduction was performed on DBD sequences using CD-HIT [40] at 40% sequence identity. The resulting non-redundant TF collection was partitioned into independent training, validation, and test sets. Ten percent of TFs were reserved as a fully independent holdout test set excluded from all stages of model optimization and architecture selection, whereas the remaining data were used in a three-fold cross-validation framework.

Nearest-neighbor TFs, using full TF sequences, were identified through all-versus-all sequence comparisons using MMseqs2 [41] and grouped into evolutionary sequence-identity intervals ranging from 30–35% to 95–100% identity as in previous works [23]. These neighbor relationships were subsequently used to derive evolutionary PWM profiles spanning a broad spectrum of sequence conservation, from distant homologs to nearly identical proteins.

### 2. PWM clustering, alignment, and tensor generation

Structure-derived PWMs generated with ModCRElib and evolutionary PWMs derived from nearest-neighbor TFs frequently differed in length, positional offset, and motif composition. To generate a unified representation suitable for machine learning, all PWMs associated with each TF were clustered and aligned prior to tensor construction.

Pairwise PWM similarity was computed using TOMTOM from the MEME Suite [42], and agglomerative clustering [43] was subsequently applied to generate coherent motif assemblies: TOMTOM p-values were transformed into adjacency-distance representations used for clustering. Two parameters controlled cluster formation: a TOMTOM significance threshold (𝑇) and a maximum agglomerative linkage distance (𝐷). Small 𝑇 values generated highly stringent clusters enriched in closely related motifs, whereas larger thresholds produced broader and more heterogeneous motif assemblies.

Within each cluster, PWMs were aligned and padded to a standardized length of 50 nucleotides, generating a common coordinate system compatible with CNN and Transformer architectures in PyTorch [44]. For training and validation datasets, only clusters containing the reference PWM, or a PWM associated with a TF sharing greater than 99% sequence identity with the reference, were retained in order to preserve biologically coherent alignments.

Importantly, relaxed clustering conditions generated noisier and more compositionally heterogeneous motif assemblies that acted as a biologically grounded form of data augmentation. Exposure to these uncertain motif configurations enabled the models to learn robust consensus representations despite ambiguity in structural and evolutionary PWM inputs.

### 3. Multi-channel tensor representation and sliding-window decomposition

Each TF was represented as a 20-channel tensor in which every channel corresponded to a PWM of dimensions 4 × 50, representing nucleotide probabilities across the aligned binding-site profile.

The first six channels contained structure-derived PWMs generated from ModCRElib structural models. Because multiple structural conformations and template-derived predictions could exist for the same TF, six PWMs were randomly sampled from the available structural pool for each tensor combination.

The remaining fourteen channels contained evolutionary PWMs derived from nearest-neighbor TFs grouped according to predefined sequence-identity intervals ranging from 30–35% to 95–100% identity. Consequently, each evolutionary channel represented motif information associated with a distinct level of evolutionary conservation relative to the query TF. When no homologous TF existed within a specific interval, the corresponding channel was represented as null and masked during model processing. This masking strategy reproduced realistic inference conditions and enabled the models to learn from incomplete evolutionary information.

Because the number of possible tensor combinations was extremely large, a stochastic sampling strategy was implemented to maximize dataset diversity while maintaining computational tractability. For each TF, 500 randomized tensor combinations were generated by sampling alternative structural and evolutionary PWM configurations while preserving the same reference PWM target. This procedure acted as a biologically meaningful form of data augmentation and improved robustness to uncertainty in both structural modeling and homolog-derived motif prediction.

To make the models independent of motif length and positional offsets, aligned tensors were decomposed into overlapping sliding windows of six nucleotides using a stride of one nucleotide. For each window position, all 20 channels were simultaneously sliced, generating localized tensors of dimensions 20 × 4 × 6. The corresponding six-position segment of the experimentally derived reference PWM was extracted as the supervised target output.

Because aligned profiles frequently contained extended null regions outside the core binding motif, windows lacking sufficient informative positions were discarded. The final dataset therefore consisted of a large collection of localized tensor windows representing multiple biologically plausible structural and evolutionary motif configurations for each TF.

### 4. Independent test construction and quality stratification

The independent holdout test set was generated using the same tensor-construction and sliding-window procedures applied during training, but without restricting tensors to clusters containing the reference PWM. Instead, structural and evolutionary PWMs were sampled from all available clusters, reproducing a realistic inference scenario in which the correct motif alignment is unknown.

To systematically evaluate robustness under varying levels of motif ambiguity, predictions were stratified into multiple cluster-quality categories defined by different combinations of clustering parameters 𝑇and 𝐷. These categories ranged from highly stringent, high-confidence motif assemblies to permissive low-confidence clusters containing substantial heterogeneity. This hierarchical quality stratification enabled direct analysis of model robustness as a function of structural uncertainty, evolutionary divergence, and motif-alignment ambiguity.

### 5. Machine-learning architectures

Three complementary machine-learning architectures were implemented for PWM reconstruction: a convolutional neural network (CNN), a Transformer-based architecture, and an interpretable regression-based baseline model (details can be found in the supplementary material).

Each training sample consisted of a tensor 𝑥 ∈ ℝ^𝐵×20×4×𝖶^, where 𝐵denotes the batch size, 20 corresponds to the PWM channels associated with each TF, 4 represents nucleotide probabilities, and 𝑊corresponds to the sliding-window length (typically six nucleotides). Missing evolutionary intervals were represented using masking tensors in order to prevent absent channels from contributing to model predictions.

The CNN architecture (WindowCNNPWM) processed the complete tensor as a multi-channel image, enabling extraction of local motif patterns and cross-channel interactions through stacked convolutional and pooling layers.

The Transformer architecture (WindowTransformerPWM) treated the 20 PWM channels as interacting tokens processed through multi-head self-attention layers. This architecture dynamically determined which structural or evolutionary PWM channels were most informative for each TF context, enabling integration of information across multiple evolutionary conservation scales while accounting for masked channels.

An interpretable regression-based model (WindowRegressionPWM) was additionally implemented. This model reconstructed PWM windows through weighted combinations of the 20 input channels, providing direct interpretation of the relative contribution of structural and evolutionary motif sources.

Hyperparameters for all architectures were optimized using Optuna [45] with Tree-structured Parzen Estimator (TPE) sampling [46]. Optimization also selected the best loss function, Mean Square Error(MSE) [47] or Huber Loss (SmoothL1) [48], between predicted and reference PWM windows. Additional optimization analyses included parameter-importance estimation, convergence monitoring, and hyperparameter-interaction visualization.

### 6. Model interpretability and occupancy-corrected attribution analysis

To investigate how the neural architectures utilized structural and evolutionary PWM sources we applied the analysis by Integrated Gradients (IG) [49]. IG provided a gradient-based attribution framework that quantified the contribution of each input feature relative to a baseline representation. However, because occupancy of the 20 PWM channels was not uniformly distributed across the dataset, some evolutionary identity intervals were substantially more represented than others. This introduced a potential attribution bias favoring highly populated channels. To reduce this effect, an occupancy-corrected IG strategy was implemented in which the mean attribution score for each channel was normalized according to the empirical occupancy probability of that channel across the dataset:

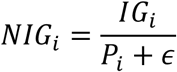

where 𝑃_𝑖_represents the fraction of samples in which channel ′𝑖′ was present and ′𝜖′ is a stabilizing constant. This normalization reduced the tendency of highly populated evolutionary intervals to dominate attribution profiles solely because of their frequency during training. Consequently, the corrected attribution scores more accurately reflected the intrinsic predictive contribution of each structural or evolutionary PWM source independently of occupancy frequency.

Interpretability analyses were performed globally across all TFs and individually for each TF, generating channel-importance profiles, positional attribution maps, TF-family-specific heatmaps, and nucleotide-level attribution distributions.

### 7. Independent evaluation using TOMTOM

Final evaluation was performed exclusively on the independent holdout TF set. Predicted PWMs were compared against experimentally derived reference motifs using TOMTOM from the MEME Suite [42]. Prediction quality was quantified using transformed TOMTOM similarity scores:

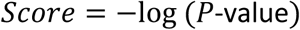

where larger values correspond to stronger agreement between predicted and reference motifs. Performance analyses were stratified according to nearest-neighbor sequence-identity intervals, TF evolutionary families, cluster-quality categories, and motif-aggregation stringencies. Summary statistics and performance distributions were visualized using boxplots and performance curves generated with Matplotlib [50].

### 8. Prediction-quality estimation and reliability modeling

An additional framework was implemented to estimate both the expected quality and reliability of each predicted PWM. The analysis used TOMTOM benchmark tables generated from the independent test set, where motif similarity was represented using transformed statistics such as −log (𝑝), −log (𝑒), and −log (𝑞) (from the statistical tests of TOMTOM yielding the P-value, E-value and Q-value, respectively) [51]. The principal response variable corresponded to the transformed TOMTOM P-value score, with larger values indicating stronger agreement between predicted and reference motifs. The effective prediction context was summarized using the available structural and evolutionary evidence from nearest-neighbor identity composition and TF-family annotation.

Three complementary strategies were used to estimate prediction quality. First, Random Forest regression models from Scikit-learn [52] were trained to predict both the expected TOMTOM similarity score and its associated dispersion using cross-validation by TF accession, thereby reducing information leakage between related samples. Because the benchmark dataset was limited in size, relatively small ensembles were used to minimize overfitting while preserving the ability to model nonlinear interactions among evolutionary similarity and TF-family-specific behavior. Second, nonlinear exponential interpolation was used to model the relationship between input-quality variables and motif-reconstruction significance. In particular, an exponential function was fitted to capture biological scenarios in which prediction quality improves with increasing evolutionary similarity or cluster consistency before progressively reaching a plateau:

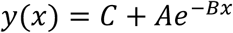

where 𝑥 represents nearest-neighbor similarity, and 𝑦(𝑥)corresponds to the transformed TOMTOM significance score.

Third, a hybrid residual-corrected framework combined the global exponential interpolation with local Random Forest refinement. Residual deviations between the observed benchmark scores and the fitted exponential trend were learned using a Random Forest regressor and subsequently added back to the baseline interpolation model. This hybrid strategy preserved the exponential formulation while improving local prediction accuracy across intermediate evolutionary-similarity regimes.

Prediction reliability was quantified by estimating uncertainty around the expected similarity score. For the Random Forest framework, reliability was derived from the predicted score dispersion, whereas for the exponential and hybrid models it was estimated using bootstrap resampling and prediction intervals. Final reliability margins were reported as the half-width of the corresponding 95% prediction interval. TF-family annotations derived from PFAM/HMMER profiles were additionally incorporated because different TF families exhibit distinct motif-conservation patterns, evolutionary constraints, and variance profiles, thereby enabling biologically informed confidence estimation for PWM reconstruction.

### 9. PWM prediction pipeline

The final PWM prediction framework reconstructs DNA-binding motifs for individual TFs or heterodimeric TF complexes starting from protein sequences and/or protein–DNA structural models. Evolutionary nearest neighbors are identified using MMseqs2, whereas supplied structures are processed through ModCRElib to derive structure-informed PWMs.

Structure-derived and evolutionary PWMs are subsequently integrated into the same 20-channel tensor representation used during benchmark construction. Candidate motifs are clustered and aligned into a common 50-position representation, after which sliding-window inference is performed using the trained CNN, Transformer, or regression architectures. Local PWM predictions generated across overlapping windows are integrated through positional averaging to reconstruct the final full-length PWM profile.

In parallel, the prediction-quality estimation framework evaluates the expected significance and reliability of each predicted PWM as a function of nearest-neighbor composition, motif-cluster quality, occupancy patterns, and TF-family context, thereby providing biologically informed confidence estimates for PWM reconstruction under varying levels of structural and evolutionary uncertainty.

Figure 1 shows the pipeline of this work. Further implementation details, clustering parameters, optimization analyses, and interpretability procedures are provided in the Supplementary Methods.

**Figure 1.**
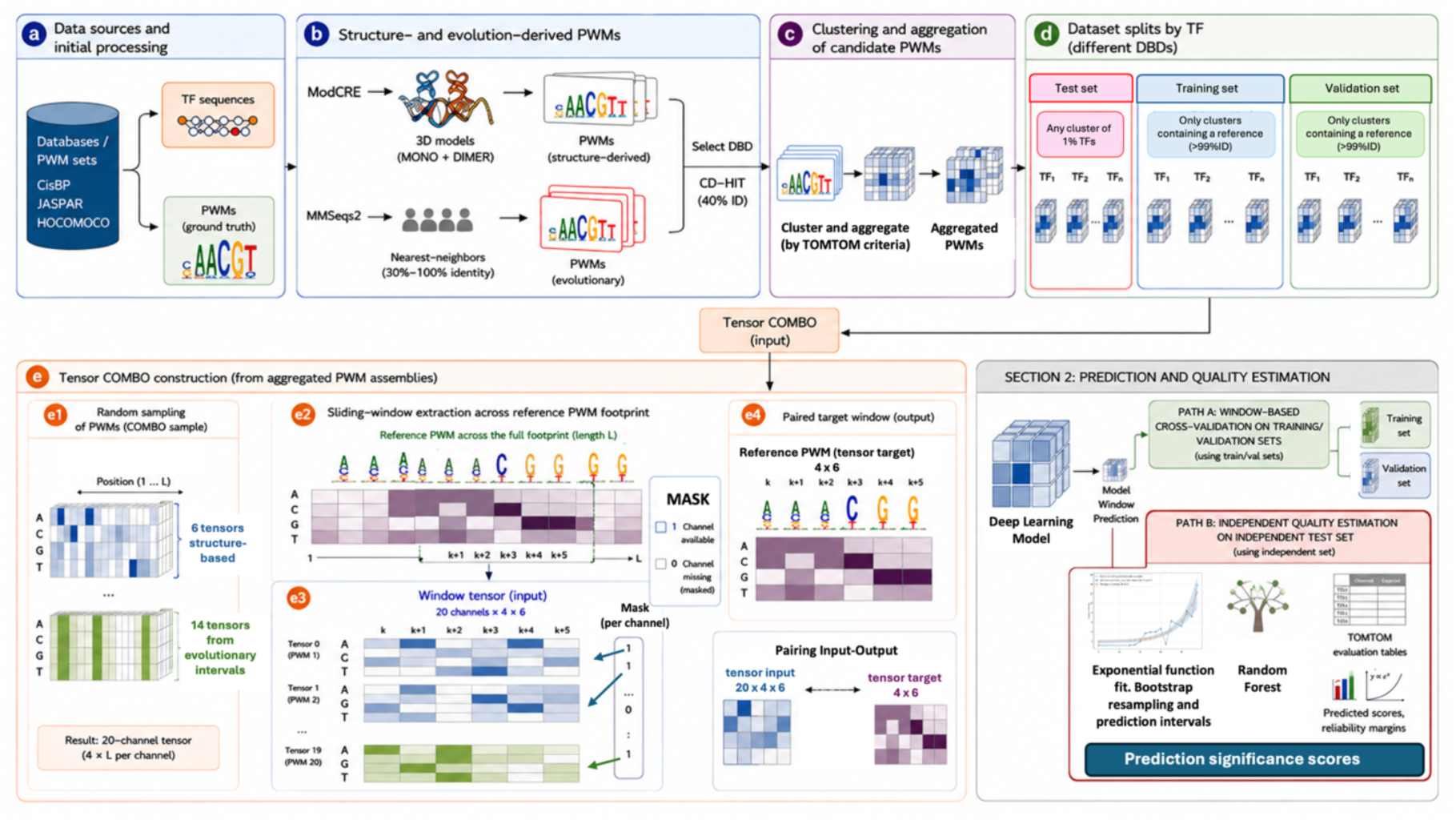
Integrated pipeline for PWM prediction and prediction-quality estimation. (a) Data sources and preprocessing. TF sequences and reference PWMs were collected from CisBP, JASPAR, and HOCOMOCO. **(b) Structure- and evolution-derived PWMs. Candidate** PWMs were generated via two complementary sources: ModCRE structural models (monomers/dimers) and MMseqs2 nearest-neighbor searches spanning 30–100% sequence identity. Non-redundant TF sets were generated using CD-HIT at 40% identity. **(c) Clustering and aggregation.** Candidate PWMs were clustered and aligned using TOMTOM similarity scores to generate aggregated PWM assemblies. **(d) Dataset partitioning**. TFs were divided into training, validation, and independent test sets based on DNA-binding domain (DBD) classes to minimize information leakage. Training and validation used clusters containing the reference PWM or >99% identity homologs, whereas the independent test set allowed any cluster configuration. **(e) Tensor COMBO construction. (e1) Random sampling.** A 20-channel tensor was generated from 6 structure-derived PWM channels and 14 evolutionary channels distributed across sequence-identity intervals. **(e2) Sliding-window extraction.** A sliding window (𝑠𝑖𝑧𝑒 = 6) traversed the aligned 50-position footprint to generate localized training examples. **(e3) Window tensor input.** The resulting 20 × 4 × 6 tensor included a channel mask indicating available (1) or missing (0) information. **(e4) Paired target output.** Each input window was paired with the corresponding 4 × 6 reference PWM window for supervised learning. **Section 2: Prediction and quality estimation.** Local windows were processed by the deep-learning framework to generate PWM predictions. **Path A: Cross-validation**. Training and validation datasets were used for model optimization and reconstruction benchmarking. **Path B: Independent quality estimation.** Independent-test predictions were compared against experimental PWMs using TOMTOM p-scores. Exponential fitting and Random Forest hybrid modeling were used to estimate the relationship between the maximum evolutionary similarity of nearest neighbors and the PWM prediction reliability. Reliability margins were estimated using bootstrap resampling. The figure was generated with ChatGPT, manually curated and refined with Gemini AI.

## RESULTS AND DISCUSSION

### 1. Systematic Hyperparameter Optimization and Training Dynamics

To maximize the predictive accuracy of the three proposed architectures (WindowRegressionPWM, WindowCNNPWM, and WindowTransformerPWM), we performed a systematic hyperparameter optimization using the Optuna framework (see methods). The optimization analysis revealed distinct parameter sensitivities for each architecture (see supplementary figure S7).

The regression-based model showed a strong dependence on the sequence window size, with an optimal value of 5, followed by weight decay regularization. The best-performing configuration was identified at Trial 14, achieving a minimum MSE of 0.2748. The CNN architecture demonstrated substantially improved predictive performance. In this model, the number of convolutional filters represented the dominant determinant of accuracy, with the optimal configuration using 128 filters and a sequence window size of 6. The best CNN configuration was identified at Trial 49 with a minimum MSE of 0.1837, although other trials achieved similar minimum and confirmed the robustness of the selected parameters. The Transformer model achieved the best overall predictive accuracy, although many trials with the same parameters were not as good as this. The embedding dimension (d_model) emerged as the most influential hyperparameter, with an optimal value of 32. The best Transformer configuration was obtained at Trial 27, reaching a minimum MSE of 0.1812.

Analysis of training dynamics revealed clear architectural differences. The regression model converged almost immediately, showing limited optimization variability across epochs. In contrast, both CNN and Transformer architectures exhibited progressive learning behavior characterized by rapid decreases in training and validation loss during the first epochs followed by stable convergence plateaus. Across all architectures, average validation losses remained consistently low, indicating robust optimization and limited overfitting.

### 2. Feature Relevance and Interpretability

We analyzed the contribution of the 20-channel input tensor using Integrated Gradients. The tensor integrates structure-derived information together with evolutionary PWMs obtained from homologous transcription factors distributed across multiple sequence identity intervals.

A preliminary analysis of tensor occupancy revealed a substantial sampling bias in the underlying biological databases. Evolutionary channels corresponding to homologs within the 30–55% sequence identity range showed the highest frequency of occurrence, together with a second pronounced enrichment for homologs within the 95–100% identity interval (see figure S8A).

Regression coefficients indicated that the strongest positive contributions originated from high-identity homolog channels, particularly those corresponding to sequence identities above 90% (Figure S8B). As we could expect, intermediate evolutionary channels also contributed positively, although with lower magnitude.

Initial Integrated Gradient analyses appeared to conflict with the learned regression weights. Raw IG values suggested that channels corresponding to the 30–55% identity interval contributed disproportionately to the final predictions (Figure S8C). We determined that this apparent discrepancy originated from the uneven occupancy of tensor channels across the dataset. Since intermediate-identity homologs are highly represented in biological databases, their high frequency artificially inflated raw attribution scores.

To correct for this effect, we implemented a Normalized Integrated Gradient (NIG) approach in which attribution scores were normalized according to channel occupancy frequency (Figure S8D). After normalization, all architectures converged toward a consistent biological interpretation (see Figure 2). In all models, Tensor 19, corresponding to the PWM derived from the closest homologs (95–100% sequence identity), emerged as the dominant predictive feature. Structure-derived tensors retained moderate but biologically meaningful contributions, accounting for approximately 25% of the importance associated with the primary feature. Evolutionary channels spanning intermediate sequence identities contributed between 30% and 45% of the total normalized importance. Importantly, the Transformer and CNN architectures showed a more balanced integration of lower-identity evolutionary channels and structure-derived tensors than the regression model. This suggests that Transformer and CNN architectures are capable of combining multiple weak evolutionary and structural signals into a refined consensus representation of DNA-binding specificity, rather than simply memorizing the PWM of the closest homolog. Structure-derived channels maintained a stable baseline relevance across all architectures, supporting the complementary contribution of structural information to the prediction of transcription factor binding specificity.

**Figure 2.**
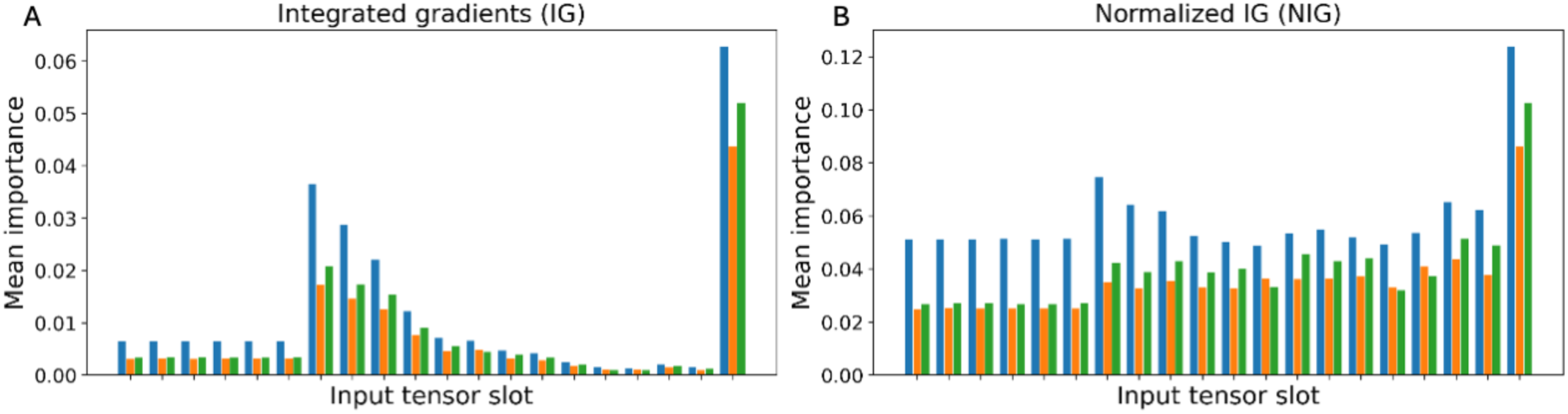
Comparative feature attribution across PWM prediction architectures. In A panel is shown the Integrated Gradient (IG) and panel B the Normalized Integrated Gradient (NIG) profiles for the WindowRegressionPWM (green), WindowCNNPWM (orange), and WindowTransformerPWM (blue) architectures.

The interpretability analysis demonstrates that PWM reconstruction is not driven exclusively by the nearest evolutionary neighbor. Although the PWM associated with the closest homolog represents the strongest predictive signal, simple models and more complex (deep neural) architectures integrate information distributed across multiple evolutionary distances together with structure-derived features.

This observation has important biological implications. It suggests that we can reconstruct transcription factor binding preferences by combining weak evolutionary evidence into a consensus representation that captures both conserved and divergent determinants of DNA-binding specificity. The contribution of structure-derived tensors further indicates that structural information provides complementary constraints capable of refining PWM prediction even when close homologous PWMs are unavailable.

Overall, the results support the idea that all architectures, and specially CNN and Transformer AI models, do not simply memorize homologous PWMs, but rather learn biologically meaningful representations integrating structural and evolutionary information across multiple levels of sequence divergence.

### 3. Comparative Evaluation of PWM Reconstruction Performance and Model Robustness

#### 3.1. Evolutionary-Distance-Dependent PWM Reconstruction Performance

To evaluate the predictive accuracy and robustness of the three proposed architectures (WindowRegressionPWM, WindowCNNPWM, and WindowTransformerPWM),reconstructed Position Weight Matrices (PWMs) were assessed on the independent holdout test set using TOMTOM similarity analysis from the MEME Suite. Prediction quality was quantified as the transformed statistical significance of the motif similarity (i.e. 𝑆𝑐𝑜𝑟𝑒 = −log (𝑝-value)), where larger values correspond to stronger agreement between predicted and experimentally derived reference motifs.

For each sequence-identity interval, we compared the AI-generated predictions against the baseline quality of the original structural and evolutionary PWMs used as model inputs. These baseline values represent the intrinsic quality of the available motif evidence prior to AI-based reconstruction.

The three architectures exhibited distinct performance regimes depending on evolutionary similarity (Figure 3). The regression-based model achieved the highest predictive performance at high sequence identities (>70–80% ID), consistent with the interpretability analyses showing that the closest homolog PWM (Tensor 19) constitutes the dominant predictive feature in the linear framework. Under these conditions, the regression model efficiently exploits highly conserved motif relationships to reconstruct accurate binding profiles.

**Figure 3.**
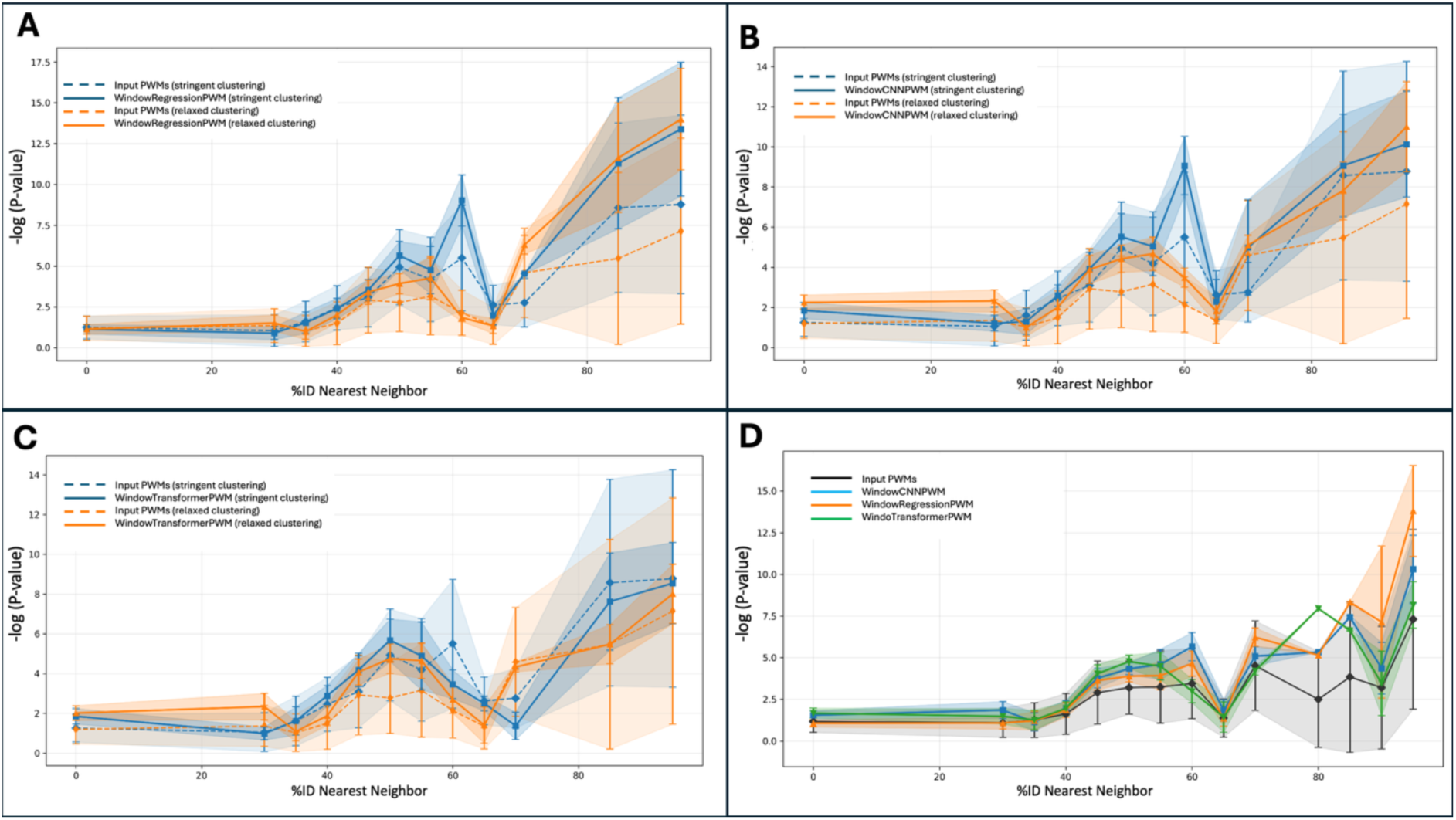
PWM reconstruction performance across evolutionary similarity intervals and clustering conditions. (A–C) Mean TOMTOM similarity scores obtained for reconstructed PWMs generated using the WindowRegressionPWM, WindowCNNPWM, and WindowTransformerPWM architectures, respectively. Scores are represented as transformed TOMTOM statistical significance values ((-\log(P\text{-value}))) as a function of the sequence identity of the closest homolog present in the 20-channel tensor input representation. Continuous lines correspond to AI-generated predictions, whereas dashed lines represent the baseline quality of the original structural and evolutionary PWMs used as model inputs. Blue curves indicate stringent motif clustering conditions (P < 0.001), whereas orange curves correspond to relaxed clustering (P < 0.05). Shaded regions represent standard deviations. (D) Global comparison integrating all clustering conditions. Black curves represent the mean baseline scores of the input PWMs, whereas orange, blue, and green curves correspond to predictions generated by the WindowRegressionPWM, WindowCNNPWM, and WindowTransformerPWM architectures, respectively. Across most evolutionary intervals, AI-generated predictions exhibit both improved TOMTOM agreement and reduced variance relative to the original motif inputs, indicating robust denoising and consensus reconstruction capabilities.

In contrast, the CNN and Transformer architectures demonstrated superior performance in low-similarity regimes and in conditions dominated by structure-derived information. These nonlinear models consistently outperformed the regression baseline when homologous PWMs were sparse, distant, or noisy, indicating an enhanced ability to integrate heterogeneous structural and evolutionary evidence into coherent consensus motif representations.

Importantly, all architectures generally improved upon the baseline input motifs across most sequence-identity intervals. The predicted PWMs not only exhibited higher average TOMTOM similarity scores, but also substantially reduced variance relative to the original structural and evolutionary inputs. This reduction in prediction dispersion indicates that the AI models effectively denoise heterogeneous motif assemblies and stabilize consensus DNA-binding representations rather than simply interpolating between homologous PWMs.

The quality of the initial PWM clustering additionally had a major impact on prediction performance. Stringent clustering conditions, based on highly significant TOMTOM similarities ((P < 0.001)), consistently produced the highest baseline and prediction scores. Under these conditions, both the input motif assemblies and the reconstructed PWMs exhibited lower variability and greater biological consistency.

Interestingly, even under relaxed clustering conditions ((P < 0.05)), where motif assemblies incorporated more heterogeneous and potentially conflicting PWMs, the CNN and regression models frequently generated predictions whose average scores exceeded the baseline values obtained from stringent clustering alone. This observation strongly suggests that the architectures learn latent consensus relationships capable of filtering noisy or partially conflicting motif evidence.

Overall, these results demonstrate that the proposed AI frameworks are not limited to nearest-neighbor interpolation. Instead, the architectures learn biologically meaningful consensus representations that integrate structural information, distant evolutionary relationships, and heterogeneous motif assemblies into robust PWM reconstructions.

#### 3.2. Influence of Motif Aggregation Quality and Structural Uncertainty

To investigate the relationship between motif-cluster quality and PWM reconstruction fidelity, prediction performance was stratified according to the stringency of the TOMTOM-based agglomerative clustering procedure used during tensor generation (Figure S9). This analysis enabled direct evaluation of model robustness under varying levels of motif heterogeneity, alignment ambiguity, and structural uncertainty.

Under highly stringent clustering conditions, clusters corresponding to intermediate sequence identities (60–80% ID) were relatively underrepresented. In these cases, many homologous PWMs were absorbed into larger high-confidence clusters associated with closely related motifs. Consequently, only a limited number of independent motif assemblies were retained within this evolutionary regime.

Relaxation of the clustering criteria substantially altered the cluster population landscape. More permissive clustering allowed the formation of additional motif assemblies spanning intermediate similarity intervals, although at the cost of increased motif heterogeneity and reduced average TOMTOM significance. This effect was particularly evident for clusters centered around 70–85% sequence identity, where relaxed clustering introduced broader compositional variability and greater alignment ambiguity.

Despite the increased uncertainty introduced by relaxed clustering, the AI architectures consistently improved the quality of the reconstructed PWMs relative to the original input motifs. Across nearly all sequence-identity bins, predicted profiles exhibited upward shifts in TOMTOM similarity distributions together with narrower dispersion margins. This trend demonstrates that the architectures effectively filter noisy motif assemblies and extract stable consensus representations from partially conflicting structural and evolutionary evidence.

The contribution of structure-derived PWMs was particularly notable in the absence of close homologous motifs. For the 0% identity interval, corresponding exclusively to structure-derived PWMs with ModCRE, the CNN and Transformer architectures generated substantially higher-quality motifs than the regression framework. Under relaxed clustering conditions, structure-based predictions reached transformed TOMTOM scores above 3, demonstrating that three-dimensional structural information alone contains sufficient signal to reconstruct biologically meaningful DNA-binding profiles even in the absence of strong evolutionary conservation.

Interestingly, the CNN and Transformer models maintained relatively stable performance across broad evolutionary ranges, suggesting that nonlinear architectures effectively combine weak evolutionary constraints distributed across multiple identity intervals. In contrast, the regression model remained strongly dependent on highly conserved homologous PWMs.

These results indicate that relaxed clustering acts as a biologically grounded form of noise augmentation. Exposure to heterogeneous motif assemblies during training appears to encourage the neural architectures to learn generalized consensus representations that remain robust under conditions of substantial structural uncertainty and evolutionary divergence.

Under stringent clustering, intermediate evolutionary intervals (60–80% ID) are poorly populated because many homologous PWMs are incorporated into larger high-confidence clusters. Relaxation of the clustering criteria increases motif heterogeneity and alignment ambiguity, reducing the average baseline TOMTOM similarity scores. Nevertheless, AI-generated predictions consistently exhibit improved reconstruction quality and reduced variance relative to the original motif inputs across most sequence-identity intervals. Structure-derived predictions (0% ID) generated by CNN and Transformer models achieve biologically meaningful TOMTOM scores even in the absence of close homologous motifs.

#### 3.3. Modeling Prediction Reliability and Expected PWM Reconstruction Quality

Beyond PWM reconstruction itself, we next investigated whether the expected quality and reliability of the predicted motifs could be estimated directly from the structural and evolutionary composition of the input tensors. To achieve this, we modeled the relationship between nearest-neighbor evolutionary similarity, motif-cluster quality, and TOMTOM reconstruction significance using empirical benchmark statistics derived from the independent test set.

Expected PWM quality was estimated using the mean TOMTOM similarity scores associated with each evolutionary regime, whereas prediction reliability was quantified using the corresponding dispersion of the benchmark distributions. The resulting curves provide an empirical framework for estimating both the expected motif quality and the associated uncertainty under different clustering and evolutionary conditions (Figure 4).

**Figure 4.**
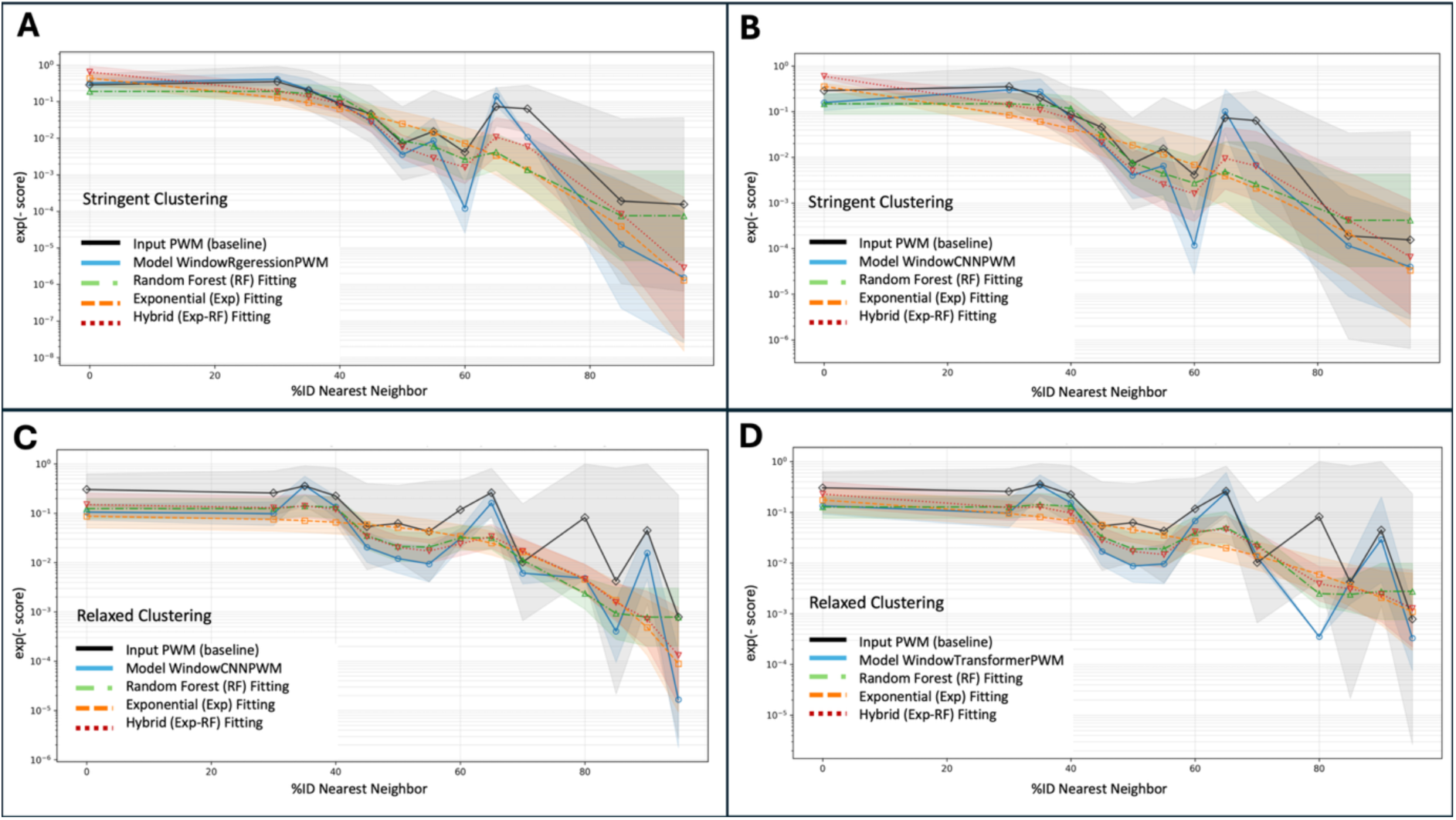
Reliability modeling and expected PWM reconstruction significance across AI architectures. Expected TOMTOM similarity scores estimated from benchmark-derived motif reconstruction statistics. Black curves represent the mean baseline quality of the original structural and evolutionary PWMs used as inputs, whereas blue curves correspond to AI-generated predictions. (A) WindowRegressionPWM predictions using stringent clustering conditions. (B–C) WindowCNNPWM predictions using stringent and relaxed clustering conditions, respectively. (D) WindowTransformerPWM predictions using relaxed clustering conditions. Prediction uncertainty is represented by shaded confidence regions corresponding to the standard deviation of the benchmark distributions. Fitting approaches estimating the P-value: exponential interpolation (orange dashed lines), Random Forest regression (green dash-dotted lines), and a hybrid residual-corrected model integrating both strategies (red dotted lines).

For each architecture, the highest-performing clustering regime was selected for reliability modeling. Regression-based predictions were best approximated using stringent motif clusters, whereas CNN and Transformer architectures achieved improved performance under more permissive clustering conditions, suggesting that nonlinear models benefit from exposure to heterogeneous motif assemblies.

Due to the limited number of benchmark observations in certain evolutionary intervals, three complementary fitting strategies were applied to model the expected TOMTOM significance curves: (i) nonlinear exponential interpolation, (ii) Random Forest regression, and (iii) a hybrid residual-corrected framework integrating both approaches. Among these methods, the hybrid model consistently provided the most stable and biologically realistic approximation of PWM reconstruction quality across evolutionary similarity ranges.

The reliability margins revealed important biological trends. Predictions based on close homologs exhibited both higher expected TOMTOM significance and lower uncertainty, whereas predictions generated from distant homologs or purely structure-derived PWMs displayed broader confidence intervals. Nevertheless, CNN and Transformer architectures maintained relatively robust performance even under sparse evolutionary conditions, particularly when relaxed motif clustering enabled integration of complementary structural evidence.

Importantly, the reliability analysis demonstrates that PWM prediction uncertainty is not random but strongly structured by evolutionary similarity, motif-cluster consistency, and transcription-factor family context. Consequently, the framework provides not only reconstructed motifs but also biologically interpretable confidence estimates associated with each prediction scenario.

Due to sparse benchmark observations in several evolutionary intervals, three complementary fitting approaches were applied to estimate the expected reconstruction quality: exponential interpolation, Random Forest regression, and a hybrid residual-corrected model integrating both strategies. The hybrid framework provided the most stable approximation of PWM reconstruction quality across evolutionary similarity regimes.

The results demonstrate that PWM reconstruction quality and reliability strongly depend on the combined effects of evolutionary similarity, motif-cluster consistency, and structural uncertainty. CNN and Transformer architectures maintain biologically meaningful predictive power even under low-homology conditions, particularly when integrating structure-derived PWM evidence.

#### 3.4. TF-Family-Specific Prediction Reliability

We next evaluated whether prediction reliability differed across transcription-factor families. Because TF families differ in DNA-recognition mechanisms, evolutionary constraints, motif conservation, and structural flexibility, family-specific reliability models may provide more informative confidence estimates than a single global model.

This analysis was performed using the CNN architecture under relaxed clustering conditions, which provided a favorable compromise between motif coverage and predictive robustness. For each TF family, expected TOMTOM similarity scores were compared between the original input profiles and the CNN-predicted PWMs. A hybrid reliability model combining exponential interpolation with Random Forest residual correction was then fitted when sufficient benchmark observations were available.

Only a subset of TF families contained enough data to support meaningful family-specific fitting (see Figure S10). The strongest support was obtained for zf-C2H2, Homeobox, and Zf-C4/nuclear receptor families, where the number and distribution of benchmark observations enabled estimation of both expected prediction quality and reliability margins. For other families, including HLH, bZIP, and Fork-head, the available data were sufficient to visualize general trends but too limited to support robust family-specific reliability modeling.

Despite these limitations, the analysis indicates that prediction uncertainty is family dependent. Families with stronger motif conservation and broader benchmark representation produced smoother and more reliable expected-score curves, whereas sparsely represented families showed wider confidence margins and less stable fitted trends. Robust family-specific fitting was most reliable for zf-C2H2, Homeobox, and Zf-C4/nuclear receptor families, where benchmark coverage was sufficient to estimate expected prediction quality and uncertainty. For HLH, bZIP, and Fork-head families, the limited number of observations allowed visualization of general trends but did not support strong conclusions about the fitted reliability functions. These results support the inclusion of TF-family information in future prediction-confidence models and highlights the need for larger family-balanced benchmark datasets.

#### 3.5. Application of ModCRE-NN to the reconstruction of the human CTCF DNA-binding motif

To illustrate the practical application of the ModCRE-NN framework, we applied the complete PWM prediction pipeline to the reconstruction of the DNA-binding specificity of the human transcription factor CTCF (CCCTC-binding factor), one of the best-characterized insulator proteins in vertebrates[53]. The amino-acid sequence of human CTCF was provided as input to the inference workflow described in Figure S6 and in the Supplementary Methods.

Evolutionary nearest neighbors were first identified using MMseqs2, while structure-derived PWMs were generated using ModCRE from available structural models obtained through AlphaFold predictions [31] and homology modeling [54]. Candidate PWMs derived from homologous TFs and structural information were subsequently clustered, aligned, and integrated into the standardized 20-channel tensor representation used throughout training and benchmarking. As during model training, multiple stochastic tensor combinations were generated by randomly sampling homolog-derived and structure-derived PWMs from the available pools, thereby reproducing the biologically heterogeneous conditions under which the neural architectures were optimized. For practical inference, up to five tensor combinations were evaluated for each prediction scenario, and Figure 5 shows representative predictions obtained from two selected tensor combinations with different levels of reliability.

**Figure 5.**
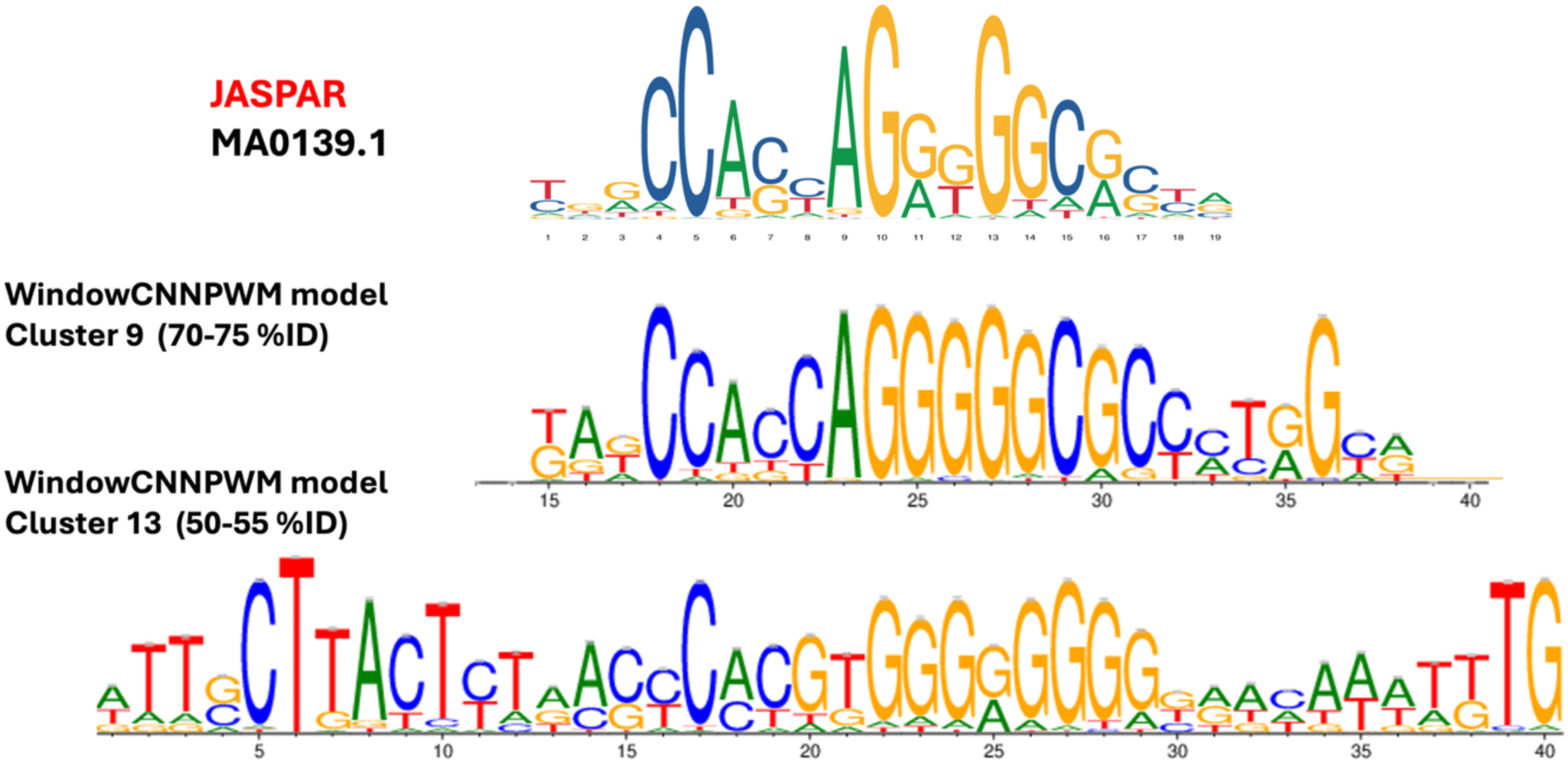
Reconstruction of CTCF DNA-binding motifs using the WindowCNNPWM architecture. Comparison between the experimental CTCF binding profile and AI-reconstructed PWMs generated under different evolutionary and structural information regimes. In the top is shown the experimental sequence logo of the human CTCF motif from the JASPAR database (MA0139.1). In the middle is shown the logo of the PWM reconstructed by the WindowCNNPWM model, using an evolutionary-dominated tensor (cluster of 13 homologs’ PWMs), with the closest homolog at 70–75% sequence identity. In the bottom is shown the logo of a PWM reconstructed using 20-channel tensors combining PWMs with evolutionary and structural information (from a cluster with 58 PWMs of homologs and 6 derived of structural models), with the closest homolog at 50–55% sequence identity.

The reconstructed motifs generated by the WindowCNNPWM architecture are shown in Figure 5. In the first prediction scenario, the prediction used an evolutionary-dominated tensor generated with PWMs of homologs clustered under relaxed conditions (𝑇 = 0.05). The input tensor was generated from one of up to five stochastic tensor combinations of 13 homologs’ PWMs, with the closest homolog located within the 70–75% sequence identity interval. No structure-derived PWMs were included in this tensor combination. We expected a significant similarity with the experimental PWM, with a P-value lower than 0.009 according to the global hybrid model and 0.0002 according to the specific family of ZF-C2H2. The CNN architecture reconstructed a motif highly similar to the canonical CTCF profile from JASPAR, indicating that the model can effectively integrate heterogeneous evolutionary evidence into a coherent consensus representation.

In the second prediction scenario, the tensor combined evolutionary PWMs together with six structure-derived PWMs generated with ModCRE from AlphaFold and homology-based structural models. In this case, the closest homolog was located only within the 50–55% sequence-identity interval, representing a substantially more challenging evolutionary regime. The expected significance of the similarity between the predicted and the experimental PWM was about 0.017 (0.006 according to the ZF-C2H2 family). Notably, the predicted motif expanded beyond the core consensus sequence and covered additional flanking specificity signals for the CTCF binding.

These example illustrates several important properties of the ModCRE-NN framework. First, PWM reconstruction is not exclusively dependent on the presence of nearly identical homologous motifs. Second, the AI models can integrate weak evolutionary constraints together with structural information to recover biologically meaningful DNA-binding profiles. Finally, the stochastic tensor-sampling strategy allows the framework to explore multiple plausible combinations of structural and evolutionary evidence, increasing robustness against uncertainty in both motif clustering and structural modeling. Together, these results demonstrate the applicability of ModCRE-NN for practical inference of TF DNA-binding specificity under heterogeneous evolutionary and structural conditions.

## CONCLUSIONS

In this work, we developed a comprehensive framework for the prediction of transcription-factor DNA-binding motifs through the integration of structural and evolutionary information using machine-learning approaches. By combining structure-derived PWMs generated from protein–DNA interaction models with homologous evolutionary motifs distributed across multiple sequence-identity intervals, we generated a unified tensor representation capable of capturing both conserved and divergent determinants of DNA-binding specificity.

The proposed framework demonstrated that PWM reconstruction can be substantially improved through the integration of heterogeneous motif evidence using AI architectures. While the regression-based model performed particularly well in high-similarity evolutionary regimes dominated by closely related homologs, CNN and Transformer architectures exhibited superior robustness under conditions of low sequence similarity, sparse evolutionary information, and increased motif uncertainty. These nonlinear models were able to integrate weak structural and evolutionary constraints into coherent consensus representations, indicating that the architectures learn biologically meaningful patterns rather than simply memorizing the PWM of the nearest homolog.

Importantly, the predicted PWMs consistently showed lower variance and improved TOMTOM similarity scores relative to the original structural and evolutionary input motifs. This observation suggests that the models effectively denoise heterogeneous motif assemblies and stabilize noisy or partially conflicting motif evidence. The ability to improve reconstruction quality even under relaxed clustering conditions demonstrates that biologically realistic uncertainty can be exploited as a form of data augmentation that enhances generalization and robustness.

Interpretability analyses further revealed that PWM reconstruction is not driven exclusively by highly conserved homologous motifs. Although the closest homologs represented the strongest predictive signal, all architectures integrated contributions from multiple evolutionary distances together with structure-derived PWMs. In particular, the Transformer architecture showed the most balanced attribution profile across structural and evolutionary channels, suggesting enhanced capacity to integrate distributed motif evidence across heterogeneous evolutionary contexts.

An additional contribution of this work is the implementation of a prediction-quality estimation framework capable of providing biologically interpretable confidence estimates for PWM reconstruction. By combining benchmark-derived TOMTOM statistics with Random Forest regression, exponential interpolation, and hybrid residual-corrected models, we were able to estimate both expected motif quality and prediction uncertainty as functions of evolutionary similarity, motif-cluster consistency, and transcription-factor family context. The results demonstrate that prediction uncertainty is highly structured and biologically informative rather than random, supporting the importance of reliability-aware PWM prediction strategies.

The family-specific analyses additionally suggest that transcription-factor evolutionary class strongly influences PWM reconstruction reliability. Families characterized by broader benchmark coverage and stronger motif conservation produced more stable and predictable reconstruction behavior, whereas sparsely represented families exhibited wider confidence intervals and greater uncertainty. These observations highlight the importance of expanding experimentally characterized TF repertoires and developing family-aware prediction frameworks in future studies.

Overall, this work demonstrates that deep-learning architectures integrating structural and evolutionary information can reconstruct biologically meaningful transcription-factor binding motifs even under conditions of substantial evolutionary divergence and structural uncertainty. Beyond PWM prediction itself, the framework provides a general strategy for integrating heterogeneous biological evidence, estimating prediction reliability, and extracting robust consensus representations from noisy molecular data.

## ACKNOWLEDGMENTS

The work was supported by Grants PID2023-150068OB-I00 and PID2024-162166NB-I00, funded by MICIUN/AEI/10.13039/501100011033 and by “ERDF A way of making Europe”, and “Unidad de Excelencia María de Maeztu” (ref: CEX2018-000792-M), as well as an FPU scholarship (ref: FPU22/02303), an SGR from the Generalitat de Catalunya (ref: 4413015318-J.SELENT/SGR-22) and Joan Oró AGAUR-FI grants from the Generalitat of Catalonia 2024 FI-1 00552 and 2024 FI-1 00227

---

**Figure S1. Construction of the non-redundant benchmark and tensor-generation workflow for TF PWM prediction.** Overview of the complete benchmark-construction strategy integrating structural and evolutionary information for transcription factor (TF) motif prediction. In a) TF sequences and experimentally derived PWMs from CisBP, JASPAR, and HOCOMOCO are used to generate non-redundant TF sets through CD-HIT clustering and nearest-neighbor identification using MMseqs2. In b) structural models of TF monomers and dimers are processed with ModCRE to generate structure-derived PWMs, whereas nearest-neighbor TFs contribute evolutionary PWMs grouped according to sequence-identity intervals. In c) candidate motifs are clustered and aggregated into aligned PWM assemblies, after which randomized 20-channel tensor combinations are generated for training, validation, and independent testing (in d). In e) the final tensor representation integrates six structure-derived PWMs and fourteen evolutionary PWM channels spanning identity intervals from 30–35% to 95–100%.

**Figure S2. Sliding-window tensor decomposition and localized PWM training strategy.** (A) **Tensor COMBO construction.** Twenty PWMs are randomly selected to generate a 20-channel tensor of dimensions 20 × 4 × 𝐿, where each channel corresponds to a 4 × 𝐿PWM representing nucleotide preferences across the theoretical DNA-binding footprint. Channels 0–5 correspond to structure-derived PWMs generated with ModCRE, whereas channels 6–19 correspond to evolutionary nearest-neighbor PWMs grouped according to sequence-identity intervals from 30–35% up to 95–100%. Missing evolutionary intervals are represented as masked channels during training and inference. (B) **Sliding-window extraction and target pairing.** A sliding window of six consecutive positions (𝑘 … 𝑘 + 5)is simultaneously extracted from all tensor channels to generate localized input tensors of dimensions 20 × 4 × 6. The same positional window is extracted from the reference PWM to generate the paired supervised target output. Only windows containing informative positions in the reference PWM are retained for training. The procedure allows CNN, Transformer, and regression models to learn localized motif patterns independently of global PWM length or positional offsets.

**Figure S3. WindowCNNPWM architecture for localized PWM reconstruction.** Architecture of the convolutional neural network (CNN) model used for PWM prediction. Input tensors consisting of 20 candidate PWMs (20 × 4 × 6) are processed through stacked convolutional blocks composed of convolution, batch normalization, ReLU activation, dropout, and pooling layers. The encoded feature representation is subsequently decoded through fully connected layers to generate predicted Position Probability Matrices (PPMs). The optimization procedure minimizes a combined loss function incorporating reconstruction accuracy (Minimum Square error) and probability normalization constraints, while prediction quality is evaluated using Pearson correlation and Jensen–Shannon divergence metrics. The schematic was generated with AI assistance (Gemini) from the CNN implementation and model configuration scripts and further manual curation.

**Figure S4. WindowTransformerPWM architecture based on multi-head self-attention.** Architecture of the Transformer-based PWM prediction model. Each candidate PWM window is flattened and projected into a latent embedding space before entering multiple Transformer encoder layers composed of multi-head self-attention and feed-forward sublayers. The model dynamically learns context-dependent weighting among structural and evolutionary PWM channels while accounting for masked inputs corresponding to missing nearest-neighbor intervals. The decoded latent representation is transformed into a predicted PWM window and optimized using reconstruction (Minimum Square Error) and normalization loss functions. The figure was generated with AI assistance (Gemini) based on the Transformer architecture and training pipeline implemented in the project scripts and further manual curation.

**Figure S5. Regression-based consensus model for interpretable PWM prediction.** Linear regression architecture used as an interpretable baseline for PWM reconstruction. The model combines the 20 candidate PWM channels through learnable weighted contributions, where the first six structure-derived channels share a common parameter and the remaining evolutionary channels receive independent coefficients. The weighted tensors are summed and reshaped into the final 4 × 6PWM prediction. This framework provides direct biological interpretability of the relative contribution of structural and evolutionary motif sources while serving as a baseline for comparison against nonlinear CNN and Transformer architectures. The schematic was generated with AI assistance (Gemini) from the regression-model implementation and further manual curation.

**Figure S6. End-to-end PWM inference and profile reconstruction workflow.** Complete prediction workflow for PWM reconstruction from structural and evolutionary evidence. Structure-derived PWMs and nearest-neighbor evolutionary PWMs are first integrated into a unified 20-channel tensor representation with masking of missing evolutionary intervals. Candidate motifs are clustered and aligned into a common 50-position theoretical binding-site representation, after which sliding-window inference is performed using trained, regression, CNN or Transformer models. Local PWM predictions generated across overlapping windows are integrated through positional averaging to reconstruct the final full-length Position Probability Matrix. The final PWM profile is subsequently converted into a Position Weight Matrix through probabilistic sequence generation and multiple-sequence alignment reconstruction. The schematic was generated with AI assistance (Gemini) based on the inference pipeline and PWM reconstruction scripts and further manual curation.

**Figure S7. Hyperparameter optimization and training dynamics across PWM prediction architectures.** (A–C) Optimization trajectories obtained from 50 Optuna trials for the WindowRegressionPWM, WindowCNNPWM, and WindowTransformerPWM architectures, respectively. Blue points represent individual optimization trials, whereas the red line indicates the best objective value identified during the optimization process. The regression model converged rapidly with limited performance variability, while CNN and Transformer architectures progressively improved toward lower MSE values. Insets show representative training and validation loss distributions across epochs for the optimal trials. CNN and Transformer models rapidly converged within the first five epochs while maintaining stable validation performance. (D) Relative hyperparameter importance across architectures. Barplot details on the importance of the parameters are shown in Figure S7.D, panels D1 (WindowRegressionPWM), D2 (WindowCNNPWM) and D3 (WindowTransformerPWM). Window size represented the dominant parameter for the regression model, whereas the number of filters and embedding dimension (d_model) were the primary determinants of predictive performance for the CNN and Transformer architectures, respectively.

**Figure S8. Tensor occupancy, regression coefficients, and attribution normalization**. (A) Global occupancy frequency of the 20-channel input tensor across the dataset. Strong enrichment is observed for homologous PWMs within the 30–55% sequence identity range and for the closest homolog interval (95–100% identity), reflecting intrinsic biases in current biological databases. (B) Regression coefficients associated with each tensor channel. Channels corresponding to high-identity homologs (>90% identity) exhibit the largest learned weights, whereas intermediate evolutionary channels retain moderate contributions. (C) Raw Integrated Gradient (IG) attribution values for the regression model. Due to occupancy imbalance, channels corresponding to highly populated intermediate-identity homologs appear artificially overrepresented. (D) Normalized Integrated Gradients (NIG) after correcting attribution scores according to feature occupancy frequency. Tensor 19, corresponding to the PWM of the closest homologs (95–100% identity), emerges as the dominant predictive feature, while structure-derived and intermediate evolutionary channels contribute moderate but biologically relevant information.

**Figure S9. Distribution of PWM reconstruction scores across clustering-quality regimes and evolutionary similarity intervals.** Boxplots showing TOMTOM similarity score distributions stratified according to the sequence identity of the closest homolog present in the motif clusters. Blue distributions correspond to the original structural and evolutionary PWM inputs (baseline), whereas orange, green, and red distributions represent predictions generated by the WindowRegressionPWM, WindowCNNPWM, and WindowTransformerPWM architectures, respectively.(A) Highly stringent clustering conditions. (B–C) Intermediate clustering stringencies. (D) Relaxed clustering conditions.

**Figure S10. TF-family-specific reliability modeling of CNN-based PWM reconstruction.** Expected TOMTOM similarity scores estimated separately for major transcription-factor families using the WindowCNNPWM model under relaxed clustering conditions. Black curves represent the mean baseline quality of the original structural and evolutionary input PWMs, whereas blue curves correspond to CNN-generated PWM predictions. Red curves show the hybrid reliability model combining exponential interpolation with Random Forest residual correction. Shaded regions indicate reliability margins associated with each curve. Panels show family-specific models for zf-C2H2 (A), HLH (B), bZIP (C), Homeobox (D), Zf-C4/nuclear receptor (E), and Fork-head (F) transcription factors.

## REFERENCES

1. Andersson R, Sandelin A: Determinants of enhancer and promoter activities of regulatory elements. Nat Rev Genet 2020, 21(2):71–87.

2. Vorontsov IE, Kozin I, Abramov S, Boytsov A, Jolma A, Albu M, Ambrosini G, Faltejskova K, Gralak AJ, Gryzunov N et al: Cross-platform motif discovery and benchmarking to explore binding specificities of poorly studied human transcription factors. Commun Biol 2025, 8(1):1545.

3. Wasserman WW, Sandelin A: Applied bioinformatics for the identification of regulatory elements. Nat Rev Genet 2004, 5(4):276–287.

4. Stormo GD, Zhao Y: Determining the specificity of protein-DNA interactions. Nat Rev Genet 2010, 11(11):751–760.

5. Stormo GD, Roy B: DNA Structure Helps Predict Protein Binding. Cell Syst 2016, 3(3):216–218.

6. Yang L, Zhou T, Dror I, Mathelier A, Wasserman WW, Gordan R, Rohs R: TFBSshape: a motif database for DNA shape features of transcription factor binding sites. Nucleic Acids Res 2014, 42(Database issue):D148–155.

7. Hallikas O, Taipale J: High-throughput assay for determining specificity and affinity of protein-DNA binding interactions. Nat Protoc 2006, 1(1):215–222.

8. Berger MF, Bulyk ML: Protein binding microarrays (PBMs) for rapid, high-throughput characterization of the sequence specificities of DNA binding proteins. Methods Mol Biol 2006, 338:245–260.

9. Isakova A, Groux R, Imbeault M, Rainer P, Alpern D, Dainese R, Ambrosini G, Trono D, Bucher P, Deplancke B: SMiLE-seq identifies binding motifs of single and dimeric transcription factors. Nat Methods 2017, 14(3):316–322.

10. Melnikov A, Murugan A, Zhang X, Tesileanu T, Wang L, Rogov P, Feizi S, Gnirke A, Callan CG, Jr., Kinney JB et al: Systematic dissection and optimization of inducible enhancers in human cells using a massively parallel reporter assay. Nat Biotechnol 2012, 30(3):271–277.

11. Ambrosini G, Dreos R, Kumar S, Bucher P: The ChIP-Seq tools and web server: a resource for analyzing ChIP-seq and other types of genomic data. BMC Genomics 2016, 17(1):938.

12. Rhee HS, Pugh BF: ChIP-exo method for identifying genomic location of DNA-binding proteins with near-single-nucleotide accuracy. Curr Protoc Mol Biol 2012, Chapter 21:Unit 21 24.

13. He Q, Johnston J, Zeitlinger J: ChIP-nexus enables improved detection of in vivo transcription factor binding footprints. Nat Biotechnol 2015, 33(4):395–401.

14. Umeyama T, Ito T: DMS-Seq for In Vivo Genome-wide Mapping of Protein-DNA Interactions and Nucleosome Centers. Cell Rep 2017, 21(1):289–300.

15. Yan J, Qiu Y, Ribeiro Dos Santos AM, Yin Y, Li YE, Vinckier N, Nariai N, Benaglio P, Raman A, Li X et al: Systematic analysis of binding of transcription factors to noncoding variants. Nature 2021, 591(7848):147–151.

16. Gryzunov N, Penzar D, Kamenets V, Vyaltsev V, Kozin I, Eliseeva IA, Nozdrin V, Vorontsov IE, Bushuev S, Strekalovskikh V et al: Inferring binding specificities of human transcription factors with the wisdom of crowds. bioRxiv 2025.

17. Weirauch MT, Cote A, Norel R, Annala M, Zhao Y, Riley TR, Saez-Rodriguez J, Cokelaer T, Vedenko A, Talukder S et al: Evaluation of methods for modeling transcription factor sequence specificity. Nat Biotechnol 2013, 31(2):126–134.

18. Weirauch MT, Yang A, Albu M, Cote AG, Montenegro-Montero A, Drewe P, Najafabadi HS, Lambert SA, Mann I, Cook K et al: Determination and inference of eukaryotic transcription factor sequence specificity. Cell 2014, 158(6):1431–1443.

19. Lambert SA, Yang AWH, Sasse A, Cowley G, Albu M, Caddick MX, Morris QD, Weirauch MT, Hughes TR: Similarity regression predicts evolution of transcription factor sequence specificity. Nat Genet 2019, 51(6):981–989.

20. Rosado-Tristani DA, Albu M, Chen X, Sasse A, Laverty KU, Ray D, Tam CL, Ernst K, Lawson LP, Morris QD et al: CisBP-RNA: a web resource for eukaryotic RNA-binding proteins and their motifs. Nucleic Acids Res 2026, 54(D1):D98–D105.

21. Liu S, Gomez-Alcala P, Leemans C, Glassford WJ, Melo LAN, Lu XJ, Mann RS, Bussemaker HJ: Predicting the DNA binding specificity of transcription factor mutants using family-level biophysically interpretable machine learning. Nucleic Acids Res 2025, 53(16).

22. Gohl P, Oliva B: SNPeBoT: A method for Predicting Transcription Factor Allele Specific Binding *doi:105281/zenodo10547232* 2024.

23. Fornes O, Meseguer A, Aguirre-Plans J, Gohl P, Bota PM, Molina-Fernandez R, Bonet J, Chinchilla-Hernandez A, Pegenaute F, Gallego O et al: Structure-based learning to predict and model protein-DNA interactions and transcription-factor co-operativity in cis-regulatory elements. NAR Genom Bioinform 2024, 6(2):lqae068.

24. Mitra R, Li J, Sagendorf JM, Jiang Y, Cohen AS, Chiu TP, Glasscock CJ, Rohs R: Geometric deep learning of protein-DNA binding specificity. Nat Methods 2024, 21(9):1674–1683.

25. Yang L, Orenstein Y, Jolma A, Yin Y, Taipale J, Shamir R, Rohs R: Transcription factor family-specific DNA shape readout revealed by quantitative specificity models. Mol Syst Biol 2017, 13(2):910.

26. Fornes O, Garcia-Garcia J, Bonet J, Oliva B: On the use of knowledge-based potentials for the evaluation of models of protein-protein, protein-DNA, and protein-RNA interactions. Adv Protein Chem Struct Biol 2014, 94:77–120.

27. Zhang Y, Silvernail I, Lin Z, Lin X: Interpretable protein-DNA interactions captured by structure-sequence optimization. Elife 2025, 14.

28. de Martin X, Oliva B, Santpere G: Recruitment of homodimeric proneural factors by conserved CAT-CAT E-boxes drives major epigenetic reconfiguration in cortical neurogenesis. Nucleic Acids Res 2024, 52(21):12895–12917.

29. Gohl P, Fornes O, Bota P, Meseguer A, Bonet J, Molina-Fernandez R, Planas-Iglesias J, Hernandez AC, Gallego O, Fernandez-Fuentes N et al: ModCRElib: A standalone package to model cis-regulatory elements. bioRxiv 2026, 2026.02.18.701901.

30. Baek M, McHugh R, Anishchenko I, Jiang H, Baker D, DiMaio F: Accurate prediction of protein-nucleic acid complexes using RoseTTAFoldNA. Nat Methods 2024, 21(1):117–121.

31. Abramson J, Adler J, Dunger J, Evans R, Green T, Pritzel A, Ronneberger O, Willmore L, Ballard AJ, Bambrick J et al: Accurate structure prediction of biomolecular interactions with AlphaFold 3. Nature 2024, 630(8016):493–500.

32. Passaro S, Corso G, Wohlwend J, Reveiz M, Thaler S, Somnath VR, Getz N, Portnoi T, Roy J, Stark H et al: Boltz-2: Towards Accurate and Efficient Binding Affinity Prediction. bioRxiv 2025.

33. Manfredi M, Vazzana G, Savojardo C, Martelli PL, Casadio R: Alpha&ESMhFolds: An Updated Web Server for the Comparison, Evaluation, and Annotation of Human AlphaFold2 and ESMFold Models. J Mol Biol 2026:169663.

34. Li J, Chiu TP, Rohs R: Predicting DNA structure using a deep learning method. Nat Commun 2024, 15(1):1243.

35. Rauluseviciute I, Riudavets-Puig R, Blanc-Mathieu R, Castro-Mondragon JA, Ferenc K, Kumar V, Lemma RB, Lucas J, Cheneby J, Baranasic D et al: JASPAR 2024: 20th anniversary of the open-access database of transcription factor binding profiles. Nucleic Acids Res 2024, 52(D1):D174–D182.

36. Vorontsov IE, Eliseeva IA, Zinkevich A, Nikonov M, Abramov S, Boytsov A, Kamenets V, Kasianova A, Kolmykov S, Yevshin IS et al: HOCOMOCO in 2024: a rebuild of the curated collection of binding models for human and mouse transcription factors. Nucleic Acids Res 2024, 52(D1):D154–D163.

37. Lambert SA, Jolma A, Campitelli LF, Das PK, Yin Y, Albu M, Chen X, Taipale J, Hughes TR, Weirauch MT: The Human Transcription Factors. Cell 2018, 172(4):650–665.

38. El-Gebali S, Mistry J, Bateman A, Eddy SR, Luciani A, Potter SC, Qureshi M, Richardson LJ, Salazar GA, Smart A et al: The Pfam protein families database in 2019. Nucleic Acids Res 2019, 47(D1):D427–D432.

39. Potter SC, Luciani A, Eddy SR, Park Y, Lopez R, Finn RD: HMMER web server: 2018 update. Nucleic Acids Res 2018, 46(W1):W200–W204.

40. Fu L, Niu B, Zhu Z, Wu S, Li W: CD-HIT: accelerated for clustering the next-generation sequencing data. Bioinformatics 2012, 28(23):3150–3152.

41. Steinegger M, Soding J: MMseqs2 enables sensitive protein sequence searching for the analysis of massive data sets. Nat Biotechnol 2017, 35(11):1026–1028.

42. Bailey TL, Johnson J, Grant CE, Noble WS: The MEME Suite. Nucleic Acids Res 2015, 43(W1):W39–49.

43. Virtanen P, Gommers R, Oliphant TE, Haberland M, Reddy T, Cournapeau D, Burovski E, Peterson P, Weckesser W, Bright J et al: SciPy 1.0: fundamental algorithms for scientific computing in Python. Nat Methods 2020, 17(3):261–272.

44. Ansel J, Yang E, He H, Gimelshein N, Jain A, Voznesensky M, Bao B, Bell P, Berard D, Burovski E et al: PyTorch 2: Faster Machine Learning Through Dynamic Python Bytecode Transformation and Graph Compilation. Proceedings of the 29th ACM International Conference on Architectural Support for Programming Languages and Operating Systems, Volume 2 2024.

45. Akiba T, Sano S, Yanase T, Ohta T, Koyama M: Optuna: A Next-generation Hyperparameter Optimization Framework. In: Proceedings of the 25th ACM SIGKDD International Conference on Knowledge Discovery & Data Mining; Anchorage, AK, USA. Association for Computing Machinery 2019: 2623–2631.

46. Bergstra J, Bardenet R, Bengio Y, Kégl B: Algorithms for hyper-parameter optimization. Advances in neural information processing systems 2011, 24.

47. Fisher RA: A mathematical Examination of the Methods of determining the Accuracy of Observation by the Mean Error, and by the Mean Square Error. Monthly Notices of the Royal Astronomical Society 1920, 80(8):758–770.

48. Huber PJ: Robust Estimation of a Location Parameter. In: Breakthroughs in Statistics: Methodology and Distribution. Edited by Kotz S, Johnson NL. New York, NY: Springer New York; 1992: 492–518.

49. Sundararajan M, Taly A, Yan Q: Axiomatic attribution for deep networks. In: Proceedings of the 34th International Conference on Machine Learning - Volume 70; Sydney, NSW, Australia. JMLR.org 2017: 3319–3328.

50. Keller BW: Mastering Matplotlib 2. x : effective data visualization techniques with Python. In., 1st edition edn. Birmingham; Mumbai: Packt,; 2018: 1 online resource (214 pages).

51. Gupta S, Stamatoyannopoulos JA, Bailey TL, Noble WS: Quantifying similarity between motifs. Genome Biol 2007, 8(2):R24.

52. Pedregosa F, Varoquaux G, Gramfort A, Michel V, Thirion B, Grisel O, Blondel M, Prettenhofer P, Weiss R, Dubourg V et al: Scikit-learn: Machine Learning in Python. Journal of Machine Learning Research 2011, 12:2825–2830.

53. Spitz F, Furlong EE: Transcription factors: from enhancer binding to developmental control. Nat Rev Genet 2012, 13(9):613–626.

54. Meseguer A, Arman F, Fornes O, Molina-Fernandez R, Bonet J, Fernandez-Fuentes N, Oliva B: On the prediction of DNA-binding preferences of C2H2-ZF domains using structural models: application on human CTCF. NAR Genom Bioinform 2020, 2(3):lqaa046.

