## supplementary text and figures for "ModCRE-NN: Interpretable Deep Learning Harnesses Structural and Evolutionary Synergy to Predict Transcription Factor Binding Specificity"

**Supplementary Methods**

### 1. Construction of Training, Validation and Test Datasets for TF PWM Prediction

#### *1.1. Data Acquisition and Structural Characterization*

The dataset construction pipeline was designed to integrate both structural and evolutionary information associated with transcription factor (TF) DNA-binding specificity. TF protein sequences and experimentally derived motif libraries were collected from the JASPAR, HOCOMOCO, and Cis-BP databases, providing the reference Position Probability Matrices (PPMs) and sequence information used throughout benchmark generation and evaluation.

Structural models of TF monomers and dimers were generated using the ModCRE framework (see figure S1a). DNA-binding domain (DBD) regions were subsequently identified and extracted from these structural models in order to focus the analysis on the protein segments directly involved in DNA recognition. In parallel, TFs were annotated into evolutionary families using Hidden Markov Model profiles from Pfam together with HMMER sequence searches. These family assignments were used exclusively for downstream stratification, interpretation, and performance analysis across TF classes.

#### *1.2. Redundancy Reduction and Evolutionary Neighborhood Mapping*

To minimize homology leakage between training and evaluation datasets, redundancy reduction was performed on all extracted DBD sequences (see figure S1b). Sequences were clustered using CD-HIT at a 40% sequence identity threshold, generating a non-redundant collection of representative TFs that served as the basis for benchmark construction. This reduction strategy was specifically applied to DBD regions rather than full-length proteins because TF DNA-binding specificity is primarily determined by the DBD sequence itself.

In parallel, an all-versus-all sequence comparison was performed using MMseqs2 to identify homologous TF neighbors across the complete dataset. Evolutionary neighbors were grouped according to predefined sequence-identity intervals ranging from 30–35% to 95–100% identity. These neighbor relationships were subsequently used to derive evolutionary PWM profiles spanning a broad spectrum of sequence conservation, from distant homologs to nearly identical proteins.

#### *1.3. Dataset Partitioning and Validation Strategy*

The curated benchmark was divided into independent training, validation, and testing subsets in order to ensure rigorous model evaluation (see figure S1d). Ten percent of the TFs were reserved as a fully independent holdout test set and were excluded from all stages of model training, hyperparameter optimization, and architecture selection.

The remaining 90% of the dataset was used within a three-fold cross-validation framework. For each fold, two-thirds of the TFs were assigned to training and one-third to validation. Because redundancy reduction was applied before partitioning, all subsets satisfied the imposed non-homology constraints, thereby preventing information leakage between training and evaluation stages.

*1.4.PWM Clustering, Alignment, and Spatial Standardization*

Predicted PWMs derived from structural and evolutionary sources frequently differed in length, positional offset, and motif composition. To generate a unified representation suitable for machine learning, all PWMs associated with each TF were standardized into a common coordinate system before tensor generation (see figure S1c).

Structural PWMs generated by ModCRE and evolutionary PWMs derived from nearest-neighbor motifs were jointly clustered using Agglomerative Clustering from the scikit-learn library. Pairwise PWM similarities were calculated using TOMTOM from the MEME Suite, which estimates motif similarity through alignment-derived statistical significance values. TOMTOM p-values were transformed into an adjacency matrix used as the clustering distance representation.

To regulate motif aggregation stringency, a threshold parameter $T$ was introduced such that all TOMTOM p-values greater than $T$ were automatically assigned the maximum distance value of 1. Consequently, only statistically significant motif relationships contributed to cluster formation. A second parameter, $D$, defined the maximum linkage distance allowed during agglomerative merging. In practice, $D$ was selected as approximately two- to five-fold larger than the corresponding $T$ value, thereby controlling the balance between cluster compactness and flexibility.

Within each cluster, PWMs were aligned and padded to a fixed length of 50 nucleotides, generating a standardized spatial representation compatible with convolutional and transformer-based architectures. For training and validation datasets, only clusters containing the reference PWM, or a PWM associated with a TF sharing greater than 99% sequence identity with the reference, were retained. This restriction ensured biologically coherent motif alignments and minimized ambiguity during supervised learning.

*1.4.a Cluster Stringency and Biologically-Grounded Noise*

The clustering parameters $T$ and $D$ directly influenced the diversity and internal consistency of PWM assemblies used during tensor generation. Small threshold values, such as $T = 0.001$ or $T = 0.005$, generated highly specific clusters composed primarily of closely related motifs. Under these stringent conditions, the resulting tensor combinations exhibited reduced positional ambiguity and greater motif consistency, generally facilitating model optimization and improving training stability.

Conversely, larger thresholds, including $T = 0.01$ and $T = 0.05$, produced broader clusters containing more distantly related PWMs. This relaxed clustering strategy introduced increased positional and compositional variability into the aligned tensors, generating noisier inputs that were intrinsically more difficult for CNN and Transformer models to interpret. Nevertheless, this variability acted as a biologically grounded form of data augmentation analogous to diffusion-style perturbation strategies, but relying exclusively on experimentally or structurally derived motif information rather than synthetic noise generation. Exposure to these heterogeneous motif assemblies encouraged the models to learn robust consensus binding representations despite substantial uncertainty in the input data.

To maximize generalization capacity, training datasets incorporated PWM clusters spanning the full range of clustering stringencies, thereby exposing the models to both high-confidence and noisy motif configurations during optimization.

#### *1.5. Multi-Channel Tensor Construction*

Each TF was represented as a stack of 20 input tensors, where each tensor corresponded to a PWM of dimensions $4 \times 50$, representing the four nucleotide probabilities across the standardized 50-position motif profile (see figure S1e).

The first six channels were populated using structure-derived PWMs generated from ModCRE models. Because multiple structural conformations and template-derived predictions could exist for the same TF, six PWMs were randomly sampled from the available structural pool for each tensor combination.

The remaining fourteen channels represented evolutionary PWMs derived from nearest-neighbor TFs grouped according to sequence-identity intervals between 30–35% and 95–100% identity. If no evolutionary neighbor existed within a specific interval, the corresponding tensor channel was marked as null and masked during model processing. This masking strategy reproduced realistic inference conditions and enabled the models to learn from incomplete input configurations.

##### *1.5.a Combinatorial Sampling and Data Augmentation*

The number of theoretically possible tensor combinations for a single TF was extremely large due to the multiplicity of structural models and evolutionary neighbors. Rather than exhaustively enumerating all possible combinations, a stochastic sampling strategy was implemented to maximize dataset diversity while maintaining computational tractability.

For every TF, 500 unique randomized tensor combinations were generated. Each combination represented a distinct arrangement of structural and evolutionary PWM inputs while preserving the same reference target PWM. This procedure acted as a biologically meaningful form of data augmentation that prevented the models from overfitting to a single structural prediction or a specific homologous motif source.

The randomized sampling strategy additionally improved robustness to uncertainty present in both structural modeling and sequence-derived motif prediction. As a result, the final dataset captured a broad distribution of biologically plausible binding configurations for each TF.

#### *1.6. Sliding Window Transformation and Localized Training Pairs*

To train models independently of motif length and positional offsets, the aligned 50-position tensors were decomposed into smaller localized regions using a sliding-window strategy (see figure S2). A window of length six nucleotides was moved across the aligned profile using a stride of one nucleotide, producing 45 potential windows per tensor combination.

For each window position, all 20 channels were simultaneously sliced, generating an input tensor of dimensions $20 \times 4 \times 6$. The corresponding six-position segment of the reference PWM was extracted as the supervised target output.

Because aligned profiles frequently contained extended null regions outside the core binding motif, a quality-filtering step was applied. Only windows containing at least five informative positions out of six in the reference PWM were retained for training and validation. This filtering procedure ensured that the models learned from biologically meaningful motif regions rather than background padding.

Following randomized tensor sampling and sliding-window decomposition, each TF generated $500 \times M$independent training examples, where $M$corresponds to the number of informative windows retained after filtering. The final dataset therefore consisted of a large collection of localized $20 \times 4 \times 6$tensor inputs paired with their corresponding $4 \times 6$target PWMs.

*1.7. Construction of the Independent Test Set*

The independent holdout test set was generated using the same tensor construction and sliding-window procedures applied to the training data, but with one critical distinction. During test-set generation, tensor selection was not restricted to clusters containing the reference PWM. Instead, structural and evolutionary PWMs were sampled from all available clusters, reproducing a realistic inference scenario in which the correct motif alignment is unknown.

To systematically evaluate robustness under varying levels of input uncertainty, predictions on the independent test set were stratified according to five predefined cluster-quality categories defined by different combinations of clustering parameters $T$and $D$. These categories ranged from highly stringent, high-confidence motif assemblies to permissive, low-confidence clusters containing substantial motif heterogeneity, defined as follows:

- **High quality (stringent):** $T = 0.001$, $D = [0.002, 0.005]$
- **High-medium (less stringent):** $T = [0.001, 0.005]$, $D = [0.002, 0.005, 0.01, 0.02]$
- **Medium quality:** $T = 0.005$, $D = [0.01, 0.02]$
- **Low-medium (less relaxed):** $T = [0.005, 0.01, 0.05]$, $D = [0.01, 0.02, 0.1, 0.2, 0.05]$
- **Low quality (relaxed):** $T = [0.01, 0.05]$, $D = [0.1, 0.02, 0.05]$

This hierarchical quality stratification enabled direct analysis of model robustness as a function of motif aggregation uncertainty and provided a biologically realistic framework for evaluating PWM reconstruction under noisy and ambiguous alignment conditions.

### 2. AI Model Architectures and Learning Strategy

To reconstruct high-resolution Position Probability Matrices (PPMs) from heterogeneous structural and evolutionary information, three complementary machine-learning architectures were implemented: a Transformer-based architecture, a convolutional neural network (CNN), and an interpretable regression-based baseline model. All architectures shared the same input representation and prediction objective, thereby enabling direct comparison of learning capacity, robustness, and interpretability.

Each training sample consisted of a tensor $x \in \mathbb{R}^{B\times20\times4\times W}$, where $B$denotes the batch size, 20 corresponds to the number of PWM channels associated with each transcription factor, 4 represents the nucleotide dimensions (A, C, G, T ), and $W$denotes the sliding-window length (usually 5 or 6), typically equal to six nucleotides. Because some evolutionary sequence-identity intervals lacked neighboring TFs, a masking tensor $m \in \mathbb{R}^{B\times20}$was used to identify absent channels and prevent them from contributing to model predictions. The output of all models was a predicted PWM window $\hat{Y} \in \mathbb{R}^{B\times4\times W}$, normalized column-wise to represent nucleotide probabilities at each position.

The 20-channel input representation integrated two biologically distinct information sources. The first six channels corresponded to structure-derived PWMs generated from ModCRE structural models, whereas the remaining fourteen channels represented evolutionary PWMs grouped according to nearest-neighbor sequence identity intervals ranging from 30–35% to 95–100%. This organization enabled the models to learn how to integrate structural evidence with evolutionary conservation across multiple similarity scales.

### *2.1. Convolutional Neural Network Architecture*

The CNN architecture, termed *WindowCNNPWM* (see figure S3), interpreted the complete input tensor as a multi-channel image of dimensions $20 \times 4 \times W$. Convolutional filters operated simultaneously across nucleotide and positional dimensions, enabling the extraction of local motif features and cross-channel interactions.

Successive convolutional and pooling layers progressively compressed the representation into a compact latent feature vector, which was subsequently decoded through an MLP into the predicted PWM window. Unlike the Transformer architecture, the CNN did not explicitly apply the candidate mask during convolutional operations. Instead, missing channels were represented as zero-valued tensors, naturally minimizing their contribution during feature extraction.

The CNN architecture was computationally efficient and particularly effective at detecting localized spatial patterns and motif correlations within aligned PWM representations. Optimized hyperparameters included the number of convolutional filters, dropout rate, optimizer type, learning rate, weight decay, and batch size.

### *2.2. Transformer Architecture*

The Transformer-based architecture, termed *WindowTransformerPWM* (see figure S4), treated the 20 PWM channels as a sequence of interacting tokens. Each $4 \times W$PWM window was flattened into a vector of size $4W$and projected into a latent embedding space of dimension $d_{model}$. These embeddings were subsequently processed through multiple Transformer encoder layers composed of multi-head self-attention and feed-forward sublayers.

The self-attention mechanism enabled the model to dynamically determine which PWM channels were most informative for a given TF context. Consequently, the Transformer could prioritize high-confidence structural motifs, suppress noisy evolutionary neighbors, or combine multiple weak signals into a coherent consensus representation. The attention mechanism followed the standard formulation:

$$\text{Attention}(Q, K, V) = \text{softmax}\left(\frac{QK^T}{\sqrt{d_k}}\right)V$$

where $Q$, $K$, and $V$represent the query, key, and value matrices, respectively, and $d_k$corresponds to the dimensionality of the key vectors.

After attention-based encoding, the candidate embeddings were pooled using a mask-aware mean operation and decoded through a multilayer perceptron (MLP) to reconstruct the final PWM prediction (first producing a position Probabilistic Matrix tensor, PPM). The masking procedure ensured that absent evolutionary channels did not participate in the attention calculations. Hyperparameters optimized for the Transformer architecture included the number of attention heads, embedding dimensionality, number of encoder layers, decoder hidden size, dropout rate, optimizer type, and learning rate.

#### *2.3. Regression-Based Consensus Model*

An interpretable regression-based baseline model, termed *WindowRegressionPWM* (see figure S5), was implemented to provide a linear reference framework for comparison against the nonlinear architectures. Rather than learning complex feature interactions, this model generated predictions through a weighted combination of the 20 input PWM channels according to:

$$\hat{Y} = \sum_{i=1}^{20} w_i X_i$$

where $X_i$represents the $i^{th}$PWM tensor and $w_i$denotes its corresponding learnable coefficient.

To reduce parameter complexity, the first six structure-derived PWM channels shared a single common weight parameter, whereas the remaining fourteen evolutionary channels were assigned independent coefficients. Following weighted summation, each PWM column was normalized to preserve probability constraints.

Although incapable of modeling nonlinear interactions between PWM sources, the regression framework provided rapid convergence, stable optimization, and direct interpretability regarding the relative contribution of structural versus evolutionary information.

### 2.4. Hyperparameter Optimization Using Optuna

All architectures underwent systematic hyperparameter optimization using the Optuna framework. Separate optimization pipelines were implemented for the Transformer, CNN, and regression models. The optimization workflow consisted of loading all tensor datasets, partitioning TFs into training, validation, and independent holdout sets, sampling hyperparameter combinations using the Tree-structured Parzen Estimator (TPE) algorithm, training candidate models for a fixed number of epochs, evaluating validation performance, and selecting the best-performing configuration.

Model optimization minimized the objective function with the minimum error (MSE) or other loss functions, such as Huber Loss (SmoothL1) or $L_{opt}$, with :

$$L_{opt} = 1 - \rho(\hat{Y}, Y)$$

where $\rho$ denotes the Pearson correlation coefficient between predicted and target PWM windows. Consequently, lower objective values corresponded to stronger agreement between predicted and reference motifs. Finally, a loss function was added to reinforce the normalization of the predicted tensor (i.e. such that the contribution of all 4 nucleotides in each position summed 1.0).

The explored search spaces included architecture-specific parameters such as the number of Transformer attention heads, embedding dimensionality, encoder depth, decoder hidden size, CNN filter number, convolutional depth, regression normalization weights, and learning rates, together with shared optimization parameters including optimizer type, dropout rate, batch size, weight decay, and loss-function selection.

### 2.5. Optimization Landscape and Parameter Analysis

Optuna additionally generated diagnostic visualizations that enabled detailed interpretation of the optimization landscape. Optimization-history plots monitored convergence behavior across trials, whereas parameter-importance analyses quantified the relative contribution of individual hyperparameters to predictive performance. Parallel-coordinate and slice plots were used to visualize parameter interactions, local parameter sensitivities, and robustness across the explored search space.

These analyses provided insight into optimization stability, parameter coupling, convergence consistency, and architecture sensitivity to specific hyperparameter choices.

### 2.6. Regression Weight Analysis and Channel Importance

The regression-based model enabled direct biological interpretation of PWM-source importance through analysis of the learned channel coefficients. After training, the weights associated with the 20 input channels were extracted and visualized in order to quantify the relative contribution of structure-derived PWMs, low-identity evolutionary neighbors, and highly conserved homologous motifs.

Large positive coefficients indicated highly informative PWM sources, whereas near-zero or negative coefficients reflected noisy or weakly informative channels. Additional analyses examined occupancy distributions across the 20 evolutionary intervals in order to determine which channels were consistently represented during training and inference.

*2.7. Model Interpretability*

To investigate how the neural architectures utilized structural and evolutionary PWM sources, two complementary interpretability approaches were applied: occlusion analysis and Integrated Gradients.

Occlusion analysis quantified the importance of individual PWM channels by systematically masking one tensor at a time and measuring the resulting decrease in prediction quality. For an input tensor $x$, the importance score associated with channel $i$was defined as:

$$S_i = F(x) - F(x \setminus x_i)$$

where $F(x)$denotes the prediction score obtained using the complete input and $F(x \setminus x_i)$ corresponds to the score obtained after removing channel $i$. Large positive values indicated channels whose removal substantially degraded prediction quality.

Integrated Gradients (IG) provided a complementary gradient-based attribution framework that quantified the contribution of each input feature relative to a baseline representation. Given an input $x$and a baseline $x'$, feature attribution was calculated according to:

$$IG_i(x) = (x_i - x'_i) \times \int_0^1 \frac{\partial F(x' + \alpha(x - x'))}{\partial x_i} d\alpha$$

where the integral was numerically approximated using discrete interpolation steps. Integrated Gradients satisfy the completeness property, ensuring that the sum of feature attributions equals the difference between the prediction obtained from the input and that obtained from the baseline.

Because the occupancy of the 20 PWM channels was not uniform across the dataset, with some evolutionary identity intervals being substantially more populated than others, an occupancy-corrected Integrated Gradients strategy was additionally implemented in order to reduce attribution bias associated with feature frequency. For each channel $i$, the mean attribution score was normalized according to the empirical occupancy probability $P_i$of the corresponding channel across the dataset:

$$NIG_i = IG_i^{adj} = \frac{IG_i}{P_i + \epsilon}$$

where $P_i$represents the fraction of samples in which channel $i$was present and $\epsilon$is a small stabilizing constant preventing numerical instabilities for sparsely populated channels. This correction reduced the tendency of highly populated evolutionary intervals to dominate the attribution profiles solely due to their increased representation during training.

Consequently, the normalized attribution scores more accurately reflected the intrinsic predictive contribution of each structural or evolutionary PWM source independently of channel frequency.

Both interpretability approaches were applied globally across all TFs and individually for each TF, generating channel-importance profiles, TF-specific heatmaps, nucleotide-level attribution distributions, and positional importance maps. Analyses were additionally stratified according to TF evolutionary families and cluster-quality categories in order to evaluate how attribution patterns changed under different levels of motif uncertainty and evolutionary divergence. These analyses enabled identification of globally informative PWM channels, TF-family-specific dependency patterns, motif positions most critical for accurate prediction, and evolutionary intervals whose contributions remained informative despite low occupancy across the dataset.

**3. Independent Test Evaluation Using TOMTOM**

Final evaluation was performed exclusively on the independent holdout TF set. Predicted PWMs were compared against experimentally derived reference motifs using TOMTOM from the MEME Suite. For each TF, TOMTOM computed the statistical significance of motif similarity between predicted and reference PWMs (upon a small dataset with the PWMs of all nearest neighbors, despite this has low statistical value it still helps to qualify the similarity). The prediction quality was quantified as:

$$\text{Score} = -\log\left(P\text{-value}\right)$$

where larger values correspond to stronger motif agreement and more accurate PWM reconstruction.

Performance analyses were stratified according to nearest-neighbor sequence identity intervals, TF evolutionary families (obtained above with HMMER and PFAM), cluster-quality categories and motif aggregation stringencies. Summary statistics, including mean values and standard deviations, were visualized using boxplots and performance curves generated with the Matplotlib library.

Together, these evaluation procedures established a biologically realistic and statistically rigorous framework for assessing AI-based PWM reconstruction under varying levels of structural uncertainty, evolutionary divergence, and motif alignment ambiguity.

**4. Prediction Quality Estimation**

An additional predictive-analysis framework was implemented to estimate both the expected similarity between predicted PWMs and their unknown experimental ground-truth motifs, as well as the reliability associated with those predictions. The objective of this framework was to infer the expected TOMTOM statistical significance that a predicted PWM would achieve relative to the true motif under different evolutionary and clustering conditions, while simultaneously quantifying the uncertainty associated with that estimation. The analysis was based on the TOMTOM benchmarking tables generated during independent test evaluation, where motif similarity was represented using transformed statistics of the form

$[-\log(p), -\log(e), -\log(q)]$. The principal response variable corresponded to either the mean or maximum transformed similarity score obtained for each prediction, typically using the $-\log(p)$component as the primary measure of PWM reconstruction quality. Prediction uncertainty was estimated from the corresponding dispersion statistics observed across TOMTOM comparisons, including the standard deviation of the transformed similarity scores.

Because the analysis operated directly on benchmark evaluation tables rather than on the original 20-channel tensors, the effective prediction context was represented using a deterministic channel-availability vector derived from the nearest-neighbor identity intervals present during tensor generation. Structure-derived channels were always considered available, whereas evolutionary channels were encoded according to the presence or absence of the corresponding sequence-identity bins. This representation enabled approximation of the effective information available to the neural models during prediction without requiring reconstruction of the original tensor representations.

Importantly, the variability of the TOMTOM similarity scores depended strongly on the quality and stringency of the PWM clusters used during tensor generation. Highly stringent clustering conditions produced compact and biologically coherent motif assemblies associated with lower variance and more stable predictions, whereas permissive clustering regimes generated broader and noisier alignments that resulted in larger dispersion of TOMTOM statistics and increased uncertainty in PWM reconstruction quality. Consequently, benchmark observations spanning the complete range of cluster-quality categories were incorporated into the framework in order to jointly model both the expected prediction significance and its associated reliability margins under varying levels of motif ambiguity and structural uncertainty.

*4.1. Random Forest-Based Prediction*

The first predictive strategy employed Random Forest regression models to learn the relationship between motif-composition conditions and PWM reconstruction quality. Two independent Random Forest models were trained: one to estimate the expected transformed similarity score,

$$\hat{s} \approx -\log(\hat{p})$$

and a second model to estimate the corresponding prediction dispersion $\hat{\sigma}$, representing the expected variability of the TOMTOM significance scores.

Because the benchmark dataset was relatively limited in size, the Random Forest framework was intentionally implemented using a relatively small number of decision trees in order to reduce overfitting while preserving generalization capacity. Model evaluation was performed using GroupKFold cross-validation with TF accession identifiers as grouping variables, thereby minimizing information leakage between related samples originating from the same transcription factor.

The Random Forest framework enabled simultaneous integration of nearest-neighbor identity composition, channel occupancy patterns, cluster-quality categories, motif-aggregation stringency, and TF-family annotations derived from PFAM/HMMER profiles. This approach was particularly effective at capturing nonlinear interactions between structural uncertainty, evolutionary similarity, and TF-family-specific motif behavior. In addition to predicting the expected TOMTOM significance, the model also estimated the expected dispersion associated with each prediction condition, thereby providing a direct estimation of prediction reliability.

#### *4.2. Exponential Interpolation*

A second complementary framework was implemented using nonlinear exponential interpolation functions designed to capture the progressive relationship between input-quality metrics and PWM reconstruction significance. In particular, exponential functions were used to represent biological scenarios in which prediction quality improves rapidly with increasing motif consistency or evolutionary similarity.

The principal interpolation model was defined as:

$$y(x) = C + Ae^{-Bx}$$

where $x$ represents a quality-related variable such as nearest-neighbor sequence identity or cluster consistency, $y(x)$corresponds to the transformed TOMTOM significance score, and $A$, $B$, and $C$are fitted model parameters.

Model parameters were estimated using nonlinear least-squares regression applied to benchmark-derived observations linking evolutionary composition, cluster-quality conditions, and TOMTOM significance scores. This formulation enabled modeling of nonlinear biological behavior that could not be adequately captured using linear interpolation approaches.

For a new prediction instance $x^*$, the fitted function generated an estimate of the expected transformed significance score:

$$\hat{y}(x^*) = f(x^*; \hat{\theta})$$

which was subsequently transformed into an estimated TOMTOM p-value according to:

$$\hat{p} = 10^{-\hat{y}}$$

or equivalently:

$$\hat{p} = e^{-\hat{s}}$$

depending on the transformation scale used during modeling.

*4.3. Hybrid Residual-Corrected Exponential Framework*

To improve prediction accuracy across intermediate evolutionary-similarity regimes, a hybrid residual-corrected interpolation strategy was additionally implemented. This framework combined the global stability and interpretability of the exponential interpolation model with the local flexibility of Random Forest regression.

First, the mean transformed TOMTOM score associated with each nearest-neighbor identity interval was modeled using the exponential interpolation function described above, generating a smooth baseline trend:

$$\hat{y}_{exp}(d) = C + Ae^{-Bd}$$

where $d$ represents the nearest-neighbor sequence identity. Residual deviations from this baseline trend were subsequently calculated as:

$$r(d) = y(d) - \hat{y}_{exp}(d)$$

where $y(d)$denotes the observed benchmark score and $\hat{y}_{exp}(d)$corresponds to the fitted exponential interpolation.

A Random Forest regressor was then trained to predict these residual components:

$$\hat{r}(d) \approx r(d)$$

thereby enabling the framework to capture local deviations, intermediate nonlinearities, and irregular behaviors that could not be adequately represented by the smooth global interpolation function alone.

The final hybrid prediction corresponded to the sum of the baseline exponential trend and the learned residual correction:

$$\hat{y}_{hyb}(d) = \hat{y}_{exp}(d) + \hat{r}(d)$$

This hybrid formulation allowed the predictive framework to preserve the exponential model while simultaneously incorporating local adaptive corrections learned from the benchmark data. Importantly, this residual-correction strategy improved prediction stability without requiring reconstruction of the original high-dimensional tensor representations.

*4. Reliability Margins and Uncertainty Estimation*

A major objective of the framework was not only to estimate expected PWM reconstruction quality, but also to quantify the reliability associated with those predictions. Reliability estimates incorporated both model uncertainty and the intrinsic biological variability observed across TFs, evolutionary distances, and clustering conditions.

Prediction uncertainty was modeled using the estimated score dispersion:

$$\hat{s} \pm k\hat{\sigma}$$

with $k = 1$by default, although larger confidence factors could be applied depending on the desired confidence level.

In the nonlinear interpolation framework, uncertainty intervals were estimated using bootstrap resampling of the benchmark data. Multiple resampled datasets were generated, the exponential and hybrid models were iteratively refitted, and empirical distributions of predicted scores were obtained for each prediction condition. Prediction intervals were preferred over simple confidence intervals because they incorporate both parameter uncertainty and residual biological variability across motif-clustering conditions and TF families.

The final reliability margin was reported as the half-width of the corresponding 95% prediction interval, thereby providing a direct estimate of the expected uncertainty associated with PWM reconstruction quality under each prediction scenario.

#### *4.5. TF-Family-Specific Reliability Modeling*

The predictive framework additionally incorporated TF-family-specific information obtained from PFAM/HMMER annotations. Different TF families exhibit distinct evolutionary constraints, motif-conservation patterns, structural flexibility, and DNA-recognition mechanisms. Consequently, family-specific benchmark distributions frequently displayed markedly different TOMTOM variance profiles and reliability behaviors.

Incorporating TF-family information enabled the framework to generate biologically informed estimates of both expected PWM reconstruction significance and associated prediction reliability as a function of evolutionary similarity. Additionally, we ran several experiments to test motif-cluster quality and channel occupancy together with TF structural class and family-specific conservation behavior. This family-aware modeling strategy substantially improved interpretation of PWM prediction confidence under biologically heterogeneous conditions and provided a more realistic estimation of expected reconstruction quality across diverse transcription-factor families.

### 5. PWM Prediction Pipeline and Prediction-Quality Estimation

The final PWM prediction framework was designed to reconstruct DNA-binding motifs for individual transcription factors or heterodimeric TF complexes starting from protein sequences and/or structural models (see figure S6). Users may provide a single protein sequence for monomeric TF prediction or two sequences for heterodimeric complexes. In addition, the workflow accepts experimentally determined or computationally predicted protein–DNA complex structures, including structures generated by methods such as AlphaFold3. All supplied structures are required to contain the complete protein monomer or protein complex together with double-stranded DNA in order to preserve the structural context necessary for DNA-binding specificity inference.

The protein sequences are first used to identify evolutionary nearest neighbors within the reference TF databases using MMseqs2. In parallel, all supplied structural models are standardized into PDB format and processed through the ModCRE framework to derive structure-informed Position Probability Matrices (PPMs) representing DNA-binding preferences associated with each structure. When multiple structures are provided, each structure contributes independently to the structural PWM pool, thereby capturing conformational variability and structural uncertainty. Optionally, additional ModCRE structural models and PWMs may also be generated directly from the query sequences and incorporated into the analysis.

The evolutionary nearest-neighbor PWMs and the structure-derived PWMs are subsequently integrated into a unified candidate motif representation identical to the tensor-generation strategy used during benchmark construction. Structural PWMs populate the first six channels of the 20-channel tensor architecture, whereas evolutionary PWMs are distributed across fourteen sequence-identity intervals ranging from 30–35% up to 95–100% identity. Missing evolutionary intervals are represented as masked channels. All candidate motifs are then clustered, aligned, and aggregated using the TOMTOM-based agglomerative clustering framework in order to generate coherent motif assemblies spanning a common 50-position binding-site coordinate system.

The aligned tensors are decomposed into overlapping six-nucleotide sliding windows that are independently evaluated by the trained Transformer, CNN, or regression models. Local PWM predictions generated for each window are subsequently integrated through overlapping positional averaging to reconstruct the final full-length PWM profile. The final Position Probability Matrix is optionally converted into a Position Weight Matrix (PWM) through probabilistic sequence generation, artificial multiple-sequence alignment reconstruction, and background-normalized log-probability transformation.

In addition to PWM prediction, the framework estimates the expected quality and reliability of each predicted motif using an empirical statistical model derived from the independent benchmark analyses. This quality-estimation framework incorporates both the evolutionary identity composition of the available nearest neighbors and the clustering-quality category associated with the aggregated PWM assemblies. Because cluster stringency directly influences alignment consistency and motif heterogeneity, highly stringent clusters generally produce more reliable predictions with lower variance, whereas permissive clustering conditions introduce larger uncertainty due to increased motif ambiguity and structural variability.

### FIGURES

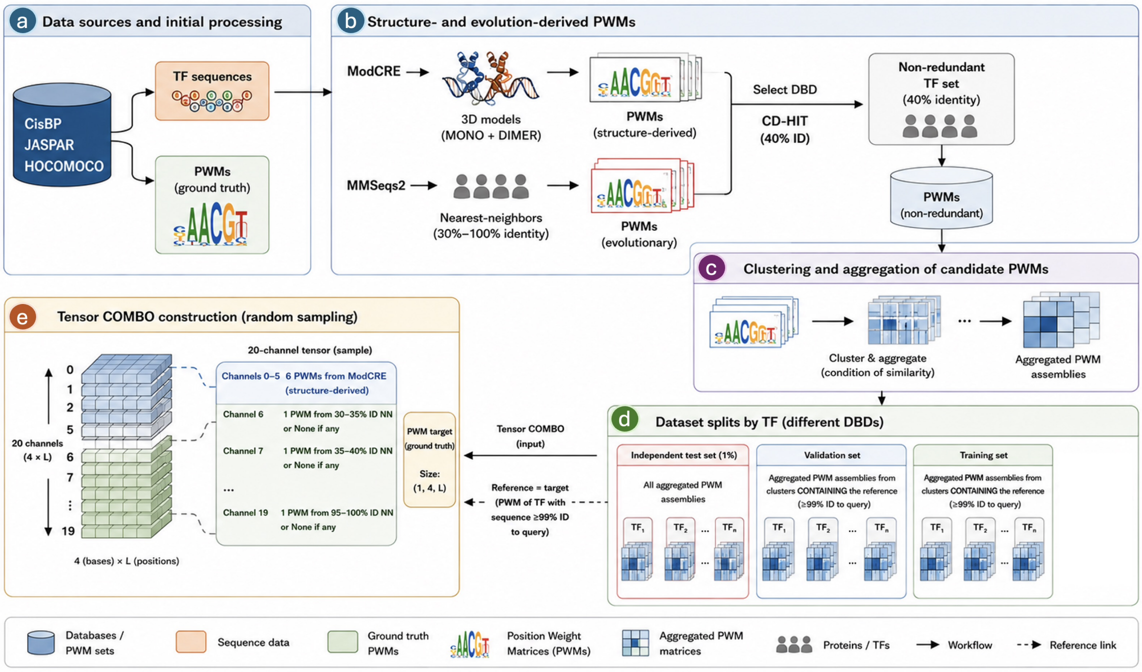


**Figure S1. Construction of the non-redundant benchmark and tensor-generation workflow for TF PWM prediction.**

Overview of the complete benchmark-construction strategy integrating structural and evolutionary information for transcription factor (TF) motif prediction. In a) TF sequences and experimentally derived PWMs from CisBP, JASPAR, and HOCOMOCO are used to generate non-redundant TF sets through CD-HIT clustering and nearest-neighbor identification using MMseqs2. In b) structural models of TF monomers and dimers are processed with ModCRE to generate structure-derived PWMs, whereas nearest-neighbor TFs contribute evolutionary PWMs grouped according to sequence-identity intervals. In c) candidate motifs are clustered and aggregated into aligned PWM assemblies, after which randomized 20-channel tensor combinations are generated for training, validation, and independent testing (in d). In e) the final tensor representation integrates six structure-derived PWMs and fourteen evolutionary PWM channels spanning identity intervals from 30–35% to 95–100%.

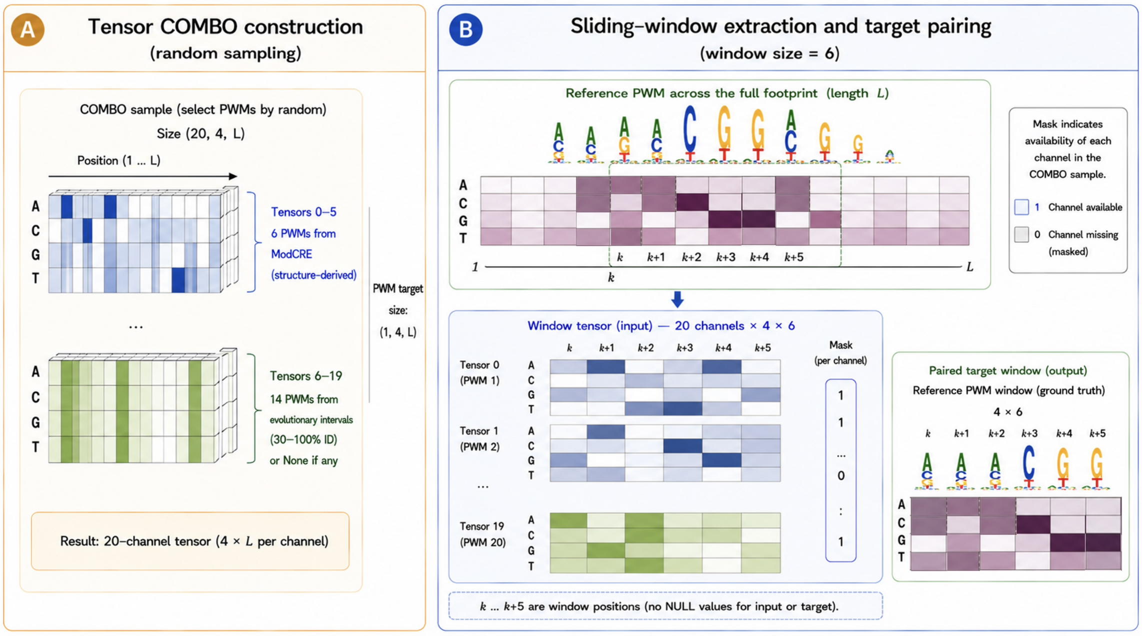


**Figure S2. Sliding-window tensor decomposition and localized PWM training strategy.**
(A) **Tensor COMBO construction.** Twenty PWMs are randomly selected to generate a 20-channel tensor of dimensions $20 \times 4 \times L$, where each channel corresponds to a $4 \times L$PWM representing nucleotide preferences across the theoretical DNA-binding footprint. Channels 0–5 correspond to structure-derived PWMs generated with ModCRE, whereas channels 6–19 correspond to evolutionary nearest-neighbor PWMs grouped according to sequence-identity intervals from 30–35% up to 95–100%. Missing evolutionary intervals are represented as masked channels during training and inference. (B) **Sliding-window extraction and target pairing.** A sliding window of six consecutive positions $(k \ldots k+5)$is simultaneously extracted from all tensor channels to generate localized input tensors of dimensions $20 \times 4 \times 6$. The same positional window is extracted from the reference PWM to generate the paired supervised target output. Only windows containing informative positions in the reference PWM are retained for training. The procedure allows CNN, Transformer, and regression models to learn localized motif patterns independently of global PWM length or positional offsets.

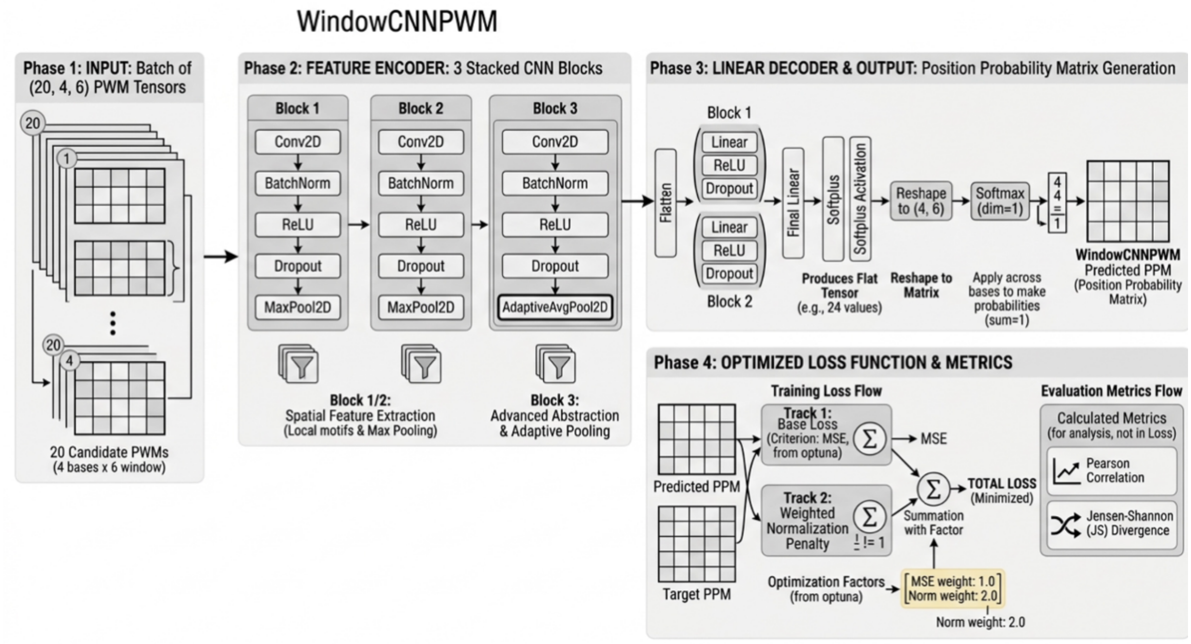


**Figure S3. WindowCNNPWM architecture for localized PWM reconstruction.**
Architecture of the convolutional neural network (CNN) model used for PWM prediction. Input tensors consisting of 20 candidate PWMs ($20 \times 4 \times 6$) are processed through stacked convolutional blocks composed of convolution, batch normalization, ReLU activation, dropout, and pooling layers. The encoded feature representation is subsequently decoded through fully connected layers to generate predicted Position Probability Matrices (PPMs). The optimization procedure minimizes a combined loss function incorporating reconstruction accuracy (Minimum Square error) and probability normalization constraints, while prediction quality is evaluated using Pearson correlation and Jensen–Shannon divergence metrics. The schematic was generated with AI assistance (Gemini) from the CNN implementation and model configuration scripts and further manual curation.

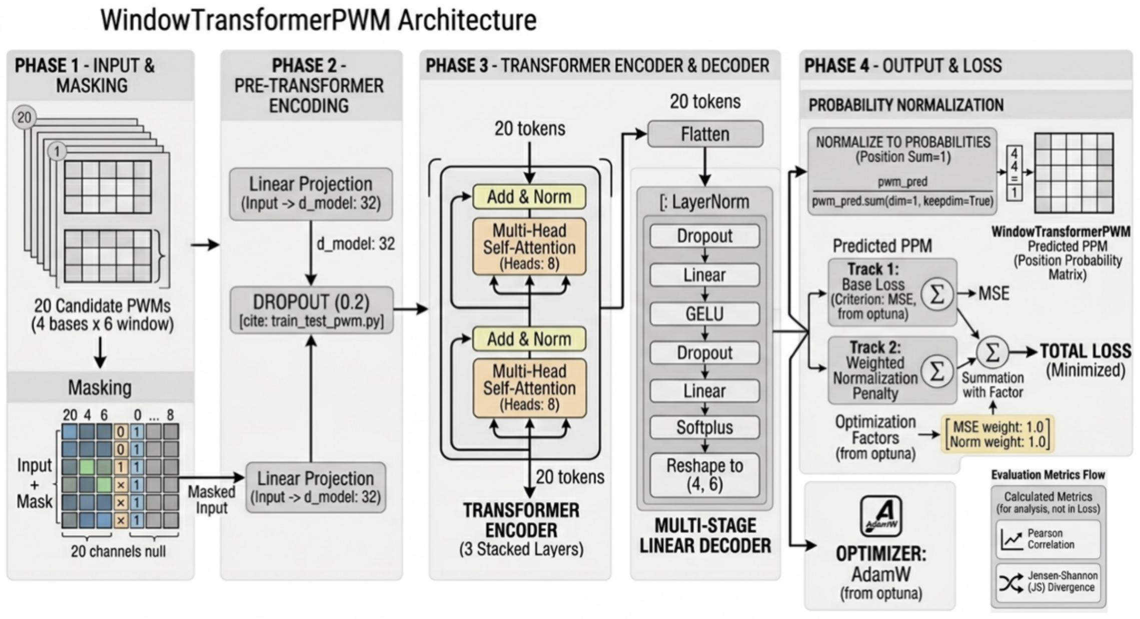


**Figure S4. WindowTransformerPWM architecture based on multi-head self-attention.** Architecture of the Transformer-based PWM prediction model. Each candidate PWM window is flattened and projected into a latent embedding space before entering multiple Transformer encoder layers composed of multi-head self-attention and feed-forward sublayers. The model dynamically learns context-dependent weighting among structural and evolutionary PWM channels while accounting for masked inputs corresponding to missing nearest-neighbor intervals. The decoded latent representation is transformed into a predicted PWM window and optimized using reconstruction (Minimum Square Error) and normalization loss functions. The figure was generated with AI assistance (Gemini) based on the Transformer architecture and training pipeline implemented in the project scripts and further manual curation.

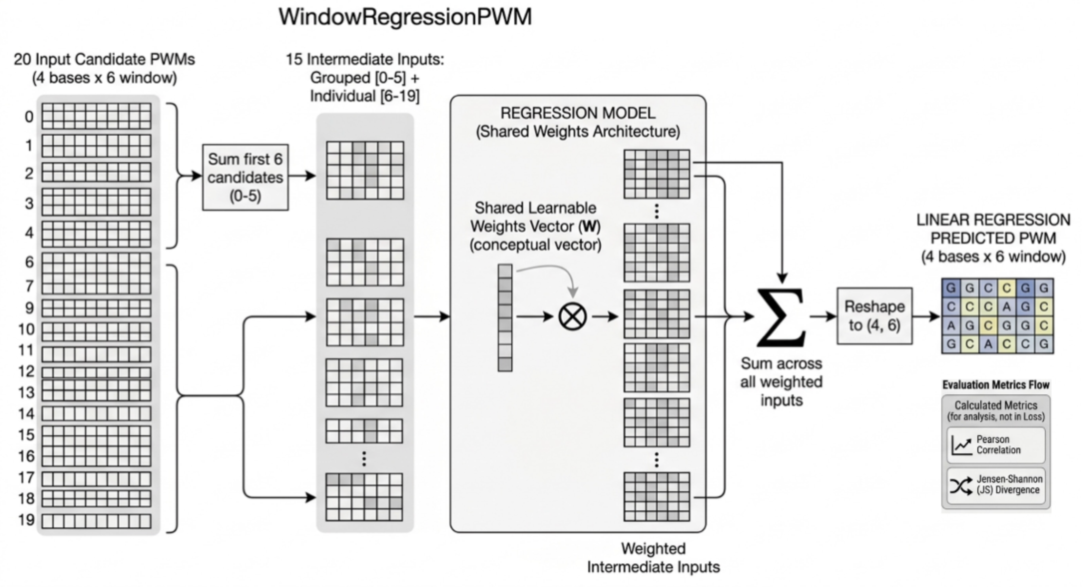


**Figure S5. Regression-based consensus model for interpretable PWM prediction.**
Linear regression architecture used as an interpretable baseline for PWM reconstruction. The model combines the 20 candidate PWM channels through learnable weighted contributions, where the first six structure-derived channels share a common parameter and the remaining evolutionary channels receive independent coefficients. The weighted tensors are summed and reshaped into the final $4 \times 6$PWM prediction. This framework provides direct biological interpretability of the relative contribution of structural and evolutionary motif sources while serving as a baseline for comparison against nonlinear CNN and Transformer architectures. The schematic was generated with AI assistance (Gemini) from the regression-model implementation and further manual curation.

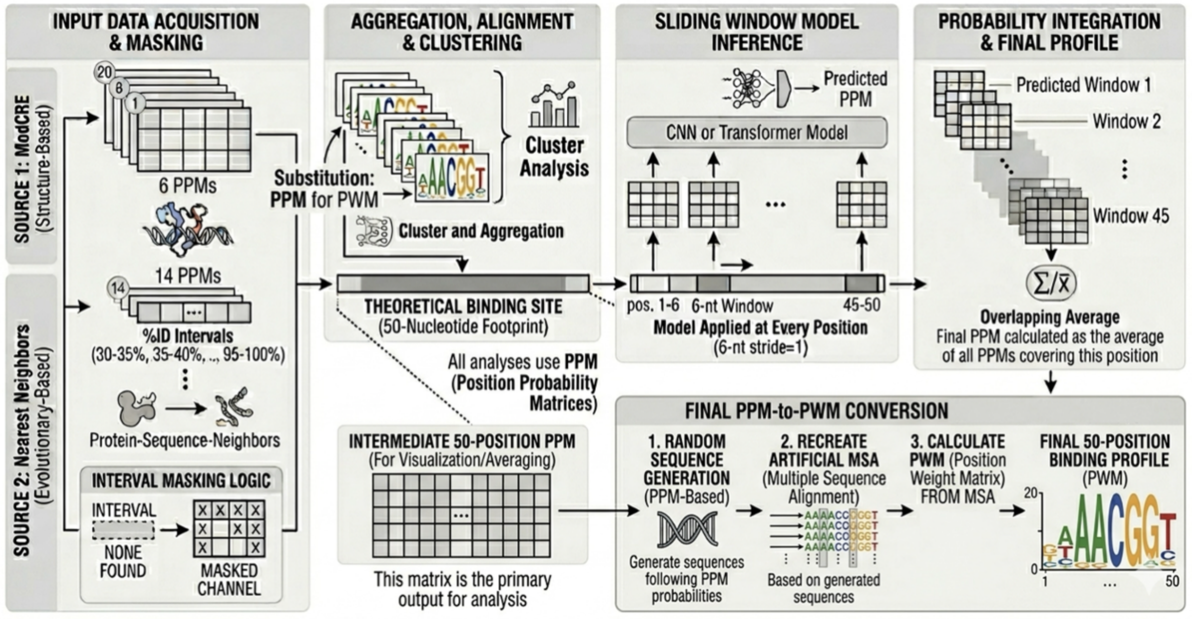


**Figure S6. End-to-end PWM inference and profile reconstruction workflow.**
Complete prediction workflow for PWM reconstruction from structural and evolutionary evidence. Structure-derived PWMs and nearest-neighbor evolutionary PWMs are first integrated into a unified 20-channel tensor representation with masking of missing evolutionary intervals. Candidate motifs are clustered and aligned into a common 50-position theoretical binding-site representation, after which sliding-window inference is performed using trained, regression, CNN or Transformer models. Local PWM predictions generated across overlapping windows are integrated through positional averaging to reconstruct the final full-length Position Probability Matrix. The final PWM profile is subsequently converted into a Position Weight Matrix through probabilistic sequence generation and multiple-sequence alignment reconstruction. The schematic was generated with AI assistance (Gemini) based on the inference pipeline and PWM reconstruction scripts and further manual curation.

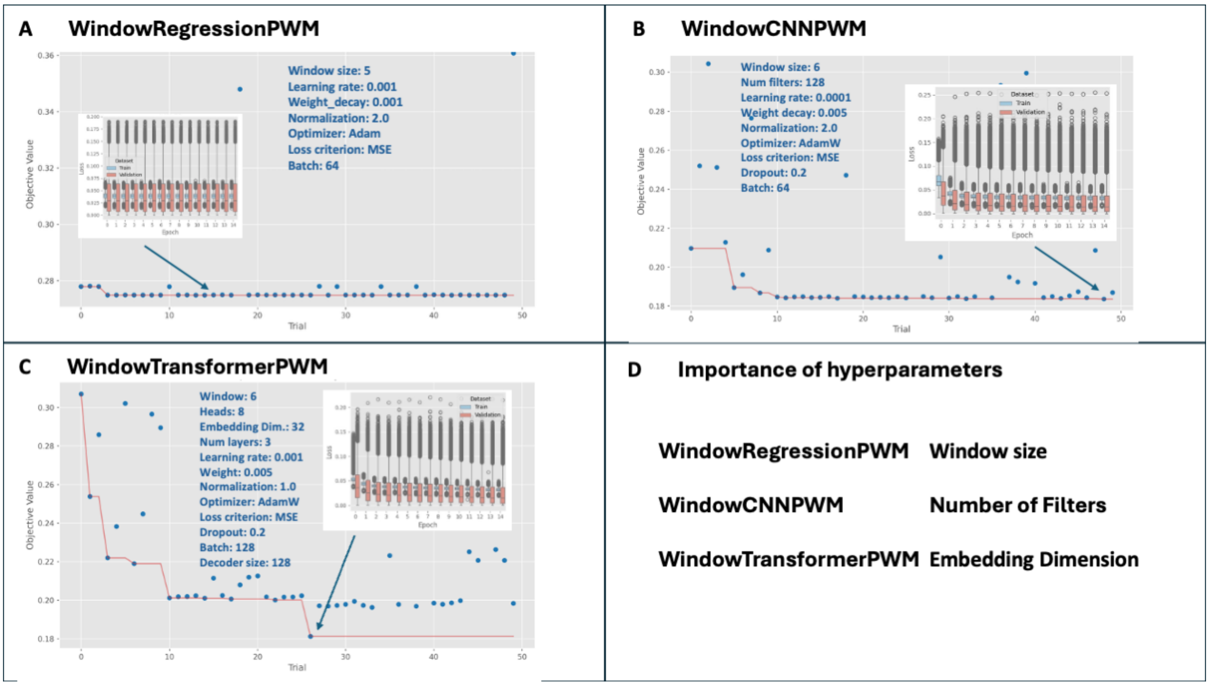


**Figure S7. Hyperparameter optimization and training dynamics across PWM prediction architectures.** (A–C) Optimization trajectories obtained from 50 Optuna trials for the WindowRegressionPWM, WindowCNNPWM, and WindowTransformerPWM architectures, respectively. Blue points represent individual optimization trials, whereas the red line indicates the best objective value identified during the optimization process. The regression model converged rapidly with limited performance variability, while CNN and Transformer architectures progressively improved toward lower MSE values. Insets show representative training and validation loss distributions across epochs for the optimal trials. CNN and Transformer models rapidly converged within the first five epochs while maintaining stable validation performance. (D) Relative hyperparameter importance across architectures. Barplot details on the importance of the parameters are shown in Figure S7.D, panels D1 (WindowRegressionPWM), D2 (WindowCNNPWM) and D3 (WindowTransformerPWM). Window size represented the dominant parameter for the regression model, whereas the number of filters and embedding dimension (d_model) were the primary determinants of predictive performance for the CNN and Transformer architectures, respectively.

S7.D1

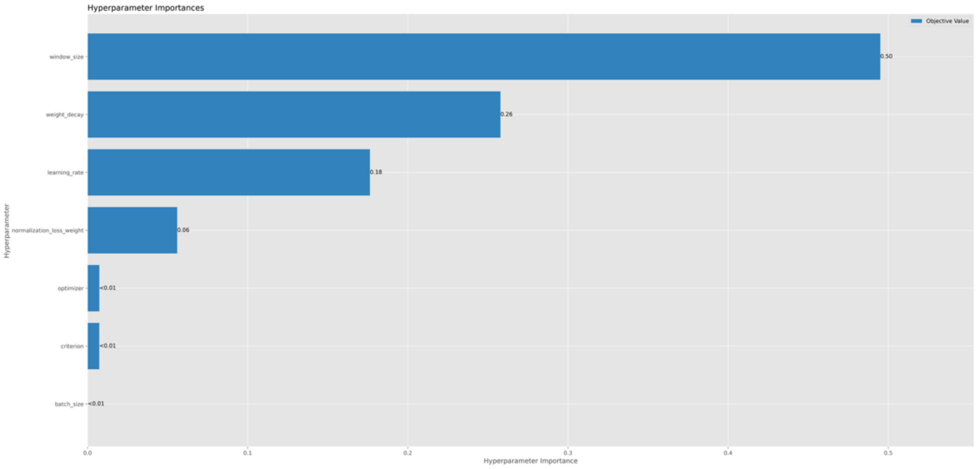


S7.D2

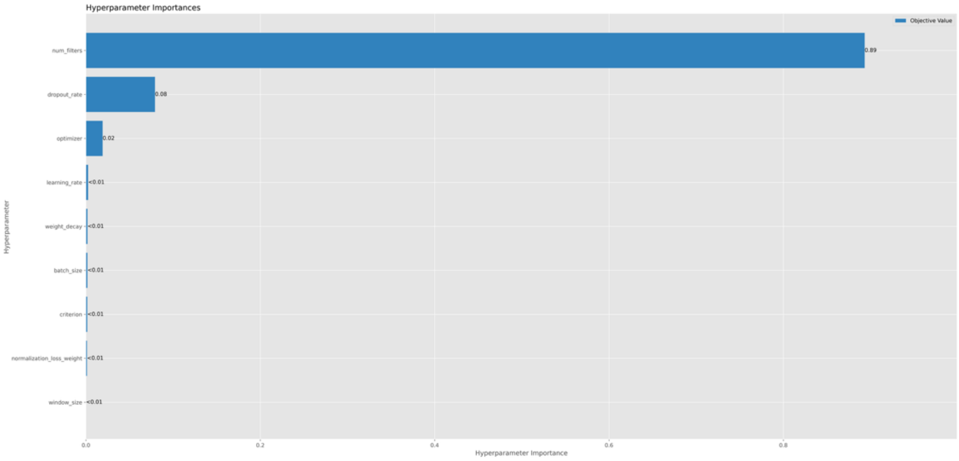


S7.D3

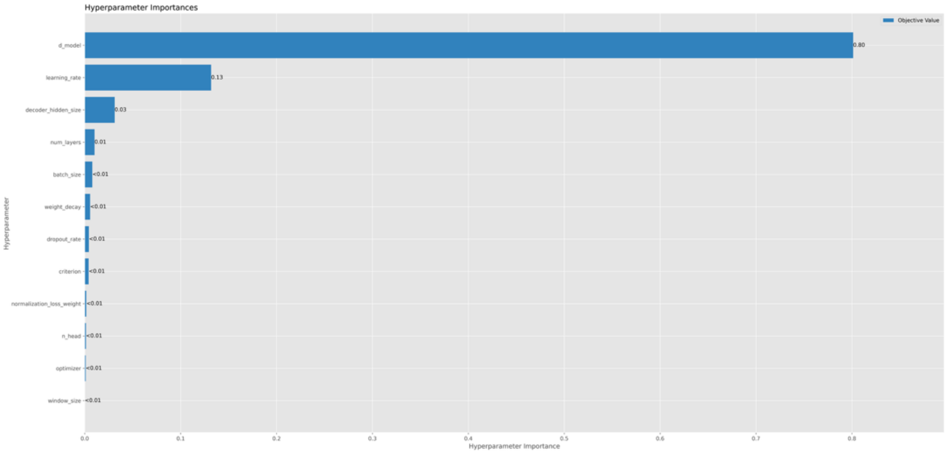


Figure S7.D (continuation)

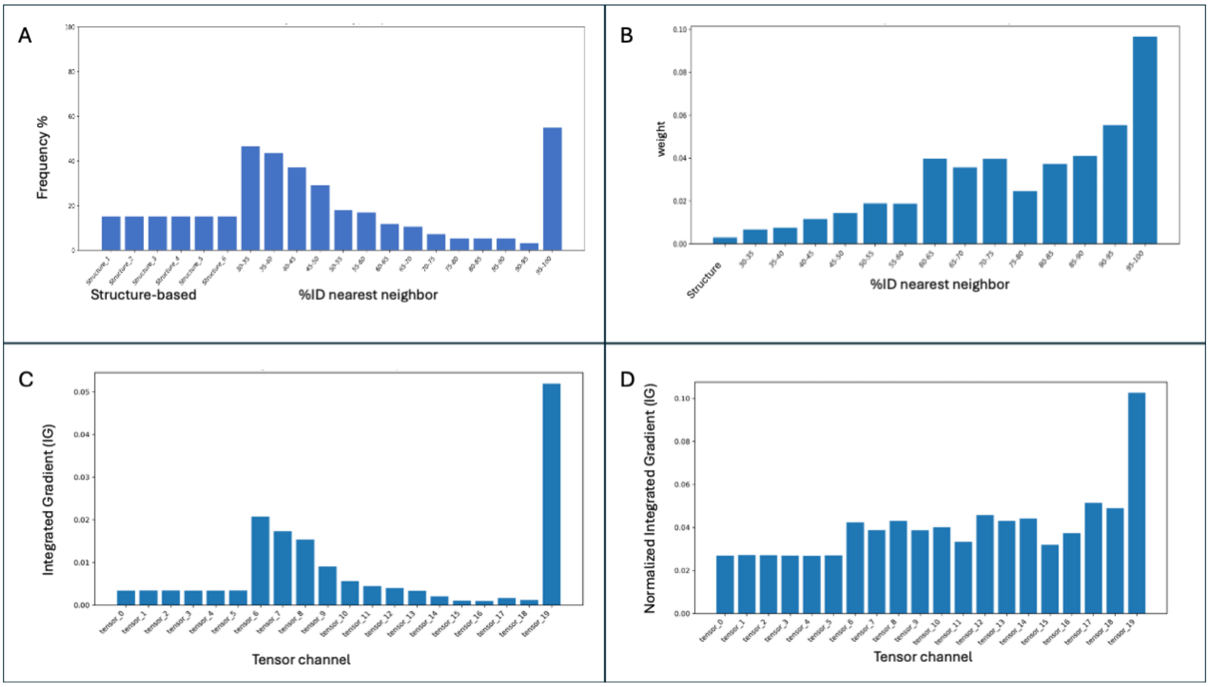


**Figure S8. Tensor occupancy, regression coefficients, and attribution normalization**. (A) Global occupancy frequency of the 20-channel input tensor across the dataset. Strong enrichment is observed for homologous PWMs within the 30–55% sequence identity range and for the closest homolog interval (95–100% identity), reflecting intrinsic biases in current biological databases. (B) Regression coefficients associated with each tensor channel. Channels corresponding to high-identity homologs (>90% identity) exhibit the largest learned weights, whereas intermediate evolutionary channels retain moderate contributions. (C) Raw Integrated Gradient (IG) attribution values for the regression model. Due to occupancy imbalance, channels corresponding to highly populated intermediate-identity homologs appear artificially overrepresented. (D) Normalized Integrated Gradients (NIG) after correcting attribution scores according to feature occupancy frequency. Tensor 19, corresponding to the PWM of the closest homologs (95–100% identity), emerges as the dominant predictive feature, while structure-derived and intermediate evolutionary channels contribute moderate but biologically relevant information.

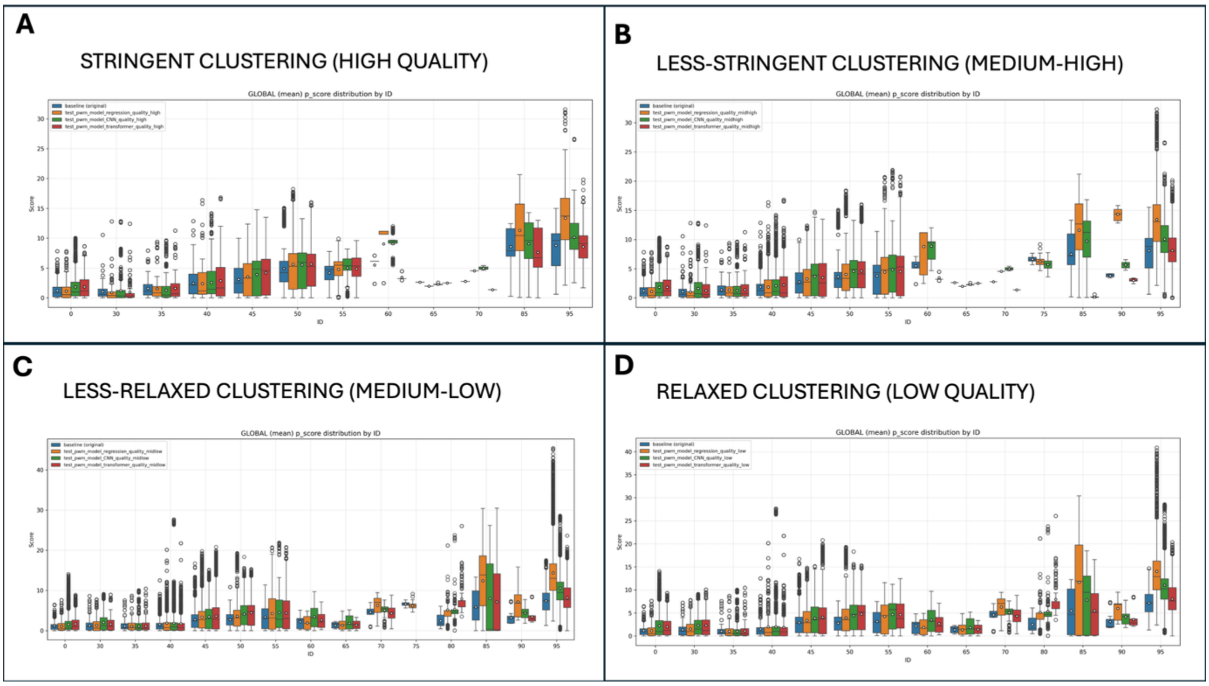


**Figure S9. Distribution of PWM reconstruction scores across clustering-quality regimes and evolutionary similarity intervals.** Boxplots showing TOMTOM similarity score distributions stratified according to the sequence identity of the closest homolog present in the motif clusters. Blue distributions correspond to the original structural and evolutionary PWM inputs (baseline), whereas orange, green, and red distributions represent predictions generated by the WindowRegressionPWM, WindowCNNPWM, and WindowTransformerPWM architectures, respectively.(A) Highly stringent clustering conditions. (B–C) Intermediate clustering stringencies. (D) Relaxed clustering conditions.

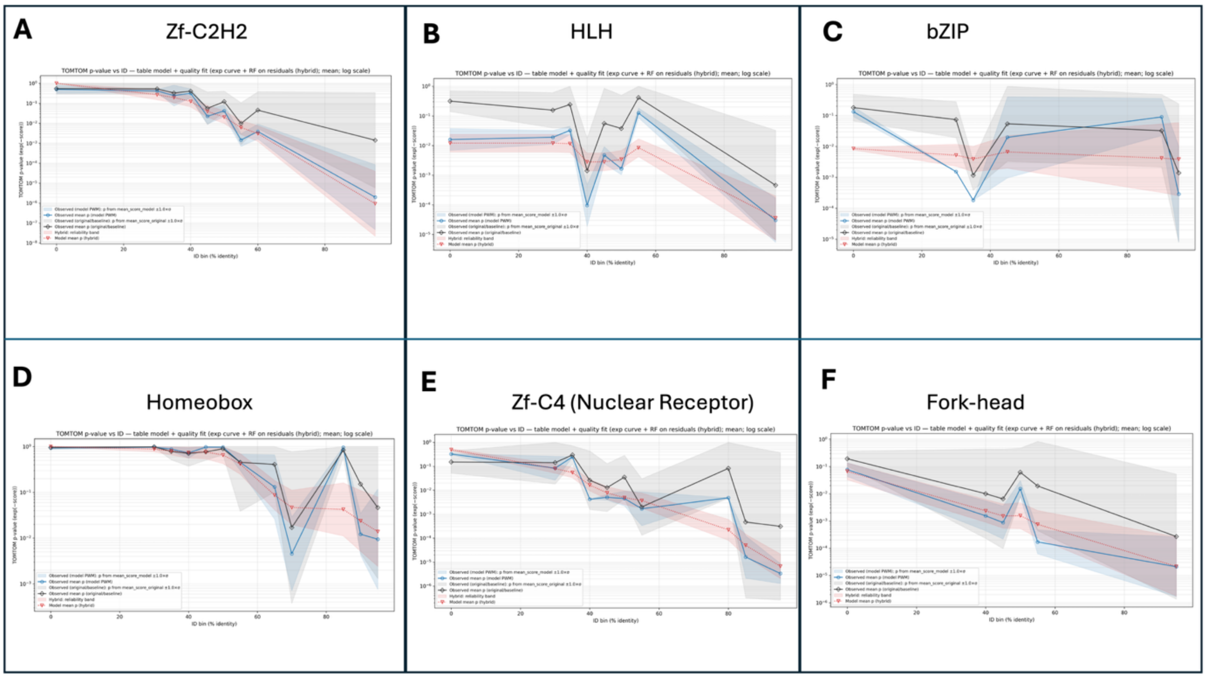


**Figure S10. TF-family-specific reliability modeling of CNN-based PWM reconstruction.** Expected TOMTOM similarity scores estimated separately for major transcription-factor families using the WindowCNNPWM model under relaxed clustering conditions. Black curves represent the mean baseline quality of the original structural and evolutionary input PWMs, whereas blue curves correspond to CNN-generated PWM predictions. Red curves show the hybrid reliability model combining exponential interpolation with Random Forest residual correction. Shaded regions indicate reliability margins associated with each curve. Panels show family-specific models for zf-C2H2 (A), HLH (B), bZIP (C), Homeobox (D), Zf-C4/nuclear receptor (E), and Fork-head (F) transcription factors.
